# Mechanism of RSL3-Induced Ferroptotic Cell Death in HT22 Cells: Crucial Role of Protein Disulfide Isomerase

**DOI:** 10.1101/2024.05.27.596002

**Authors:** Ming-Jie Hou, Xuanqi Huang, Bao Ting Zhu

## Abstract

Protein disulfide isomerase (PDI) was recently shown to be an upstream mediator of erastin-induced, glutathione depletion-associated ferroptosis through its catalysis of nitric oxide synthase (NOS) dimerization and nitric oxide (NO) accumulation. A recent study reported that RSL3, a known ferroptosis inducer and glutathione peroxidase 4 (GPX4) inhibitor, can inhibit thioredoxin reductase 1 (TrxR1). The present study seeks to test a hypothesis that RSL3 may, through its inhibition of TrxR1, facilitate PDI activation (*i.e.*, in a catalytically-active, oxidized state), thereby enhancing RSL3-induced ferroptosis through NOS dimerization and NO accumulation. Using the HT22 mouse neuronal cells as an *in-vitro* model, we show that treatment of these cells with RSL3 can strongly increase NOS protein level, and the PDI-mediated NOS dimerization is activated by RSL3, resulting in NO accumulation. Mechanistically, we find that PDI is activated in cells treated with RSL3 resulting from its inhibition of TrxR1, and the activated PDI then catalyzes NOS dimerization, which is followed by accumulation of cellular NO, ROS and lipid-ROS, and ultimately ferroptotic cell death. Genetic or pharmacological inhibition of PDI or TrxR1 partially abrogates RSL3-induced NOS activation and the subsequent accumulation of cellular NO, ROS/lipid-ROS, and ultimately ferroptosis in HT22 cells. The results of this study clearly show that PDI activation resulting from RSL3 inhibition of the TrxR1 activity contributes crucially to RSL3-induced ferroptosis in a cell culture model through the PDIl7NOSl7NOl7ROS/lipid-ROS pathway, in addition to its known inhibition of the GPX4 activity.

## INTRODUCTION

Ferroptosis is a form of oxidative cell death [1], and is morphologically distinct from apoptosis-associated characteristics [1, 2]. Mechanistically, ferroptosis can be selectively induced through cellular glutathione (GSH) depletion and/or through suppression of glutathione peroxidase 4 (GPX4), an enzyme involved in the reduction of lipid peroxides. RSL3 is a prototypical inducer of ferroptosis [3], and over the past few years, it has been widely thought that RSL3 induces ferroptosis primarily through its suppression of GPX4 function, which then leads to accumulation of cellular lipid reactive oxygen species (lipid-ROS) [4]. However, a recent study has reported that RSL3 can also strongly inhibit the catalytic activity of thioredoxin reductase 1 (TrxR1) [5]. Because TrxR1 uses NADPH as a cofactor to reduce the active-site disulfide bonds in thioredoxins, when TrxR1 is inhibited by an inhibitor (such as RSL3), the active site of the thioredoxins would stay in the oxidized state (*i.e.*, containing a disulfide bond in their active sites) [6].

Protein disulfide isomerase (PDI or PDIA1) is the prototype of the PDI family proteins, which belong to the ubiquitous dithiol/disulfide oxidoreductases of the thioredoxin superfamily [7–9]. PDI is primarily localized in the endoplasmic reticulum of mammalian cells, although a small fraction of this protein is also found in the nucleus, cytosol, mitochondria, plasma membrane and extracellular space [10]. PDI is involved in protein processing by catalyzing the formation of intra- and inter-molecular disulfide bridges in proteins [11]. Our recent studies have shown that PDI plays an important role in mediating glutamate- and erastin-induced, GSH depletion-associated oxidative cytotoxicity through PDI-mediated NOS activation (*i.e.*, homodimer formation through a disulfide bond linkage) which is followed by the accumulation of cellular NO, ROS and lipid-ROS, and ultimately the induction of ferroptotic cell death [12].

In the present study, we sought to examine the role of PDI in mediating RSL3- induced ferroptosis in HT22 neuronal cells and the mechanism of PDI activation. We found that PDI plays a crucial role in mediating RSL3-induced ferroptotic cell death in these cells through its ability to catalyze NOS dimerization, which is followed by cellular accumulation of NO, ROS and lipid-ROS, and ultimately ferroptotic cell death. Genetic or pharmacological inhibition of PDI’s function could abrogate RSL3-induced ferroptosis. The mechanism by which PDI is activated in RSL3-treated cells is caused by TrxR1 inhibition by RSL3, and this inhibition leaves more PDI proteins in the oxidized state, which is the active form to catalyze the dimerization of NOS (including both nNOS and iNOS) in HT22 cells.

## MATERIALS AND METHODS

### Reagents

RSL3 (#SML2234) and *S*-nitroso-*N*-acetylpenicillamine (SNAP, #N3398) were purchased from Sigma-Aldrich (St. Louis, MO, USA). CPZ (chlorpromazine, #S5749), ferrostatin-1 (Fer-1, #S7243), cystamine (#S3695) and 3-(4,5-dimethylthiazol-2-yl)- 2,5-diphenyltetrazolium bromide (MTT, #S6821) were purchased from Selleck Chemicals (Houston, TX, USA). Diaminofluorescein-FM diacetate (DAF-FM-DA, #S0019) and diaminofluorescein-2 diacetate (DCF-DA, #S0033) were obtained from Beyotime Biotechnology (Shanghai, China), and C11-BODIPY-581/591 (#D3861) from Thermo Fishers (Waltham, MA, USA). Anti-PDI (#3501), anti-TrxR1 (#6925) and anti-β-actin (#4970) antibodies were purchased from Cell Signaling Technology (Danvers, MA, USA); anti-GPX4 (#ab125066), anti-nNOS (#ab1376) and anti-iNOS (#ab3523) antibodies from Abcam (Waltham, MA, USA). TrxR1-siRNAs (#sc-36751) and iNOS-siRNAs (#sc- 36092) were obtained from Santa Cruz Biotechnology (Dallas, TX, USA); PDI-siRNAs and nNOS-siRNAs were from Ruibo Biotechnology (Guangdong, China). The siRNA sequences targeting the mouse TrxR1 (siTrxR1) are UUCAAGUAGAUUAGCCAAGtt (#1), UACGUUAUGAACAGCUUCCtt (#2), and UUACGUUAUGAACAGCUUCtt (#3). The siRNA sequences targeting the mouse iNOS (si-iNOS) are UUUCCUUUGUUACAGCUUCtt (#1), UUCAUGAUAACGUUUCUGGtt (#2), and UAGUAGUCCACAAUAGUACtt (#3). The siRNA sequences targeting the mouse PDI (siPDI) are CCAAGTACCAGCTGGACAA (#1), GAACGGTCATTGATTACAA (#2), and TGCTAAGATGGACTCAACA (#3). The siRNA sequences targeting the mouse nNOS (sinNOS) are GCTGCCATCCCATCACATA (#1), CCTCGTGAATGCACTCATT (#2), and GCGACAATTTGACATCCAA (#3). The siRNA sequences for negative control (siCon) are AUCCGCGCGAUAGUACGUAtt (#1), UUACGCGUAGCGUAAUACGtt (#2), and UAUUCGCGCGUAUAGCGGUtt (#3). The MDA (malondialdehyde) Assay Kit (#M496) was purchased from Dojindo Molecular Technologies (Kumamoto, Japan), and the TrxR1 Activity Assay Kit (#BC1155) from Solarbio (Beijing, China).

### Cell culture and cell viability assay

HT22 cells were obtained from the Cell Bank of the Chinese Academy of Sciences (Shanghai, China). The HT22 murine hippocampal neuronal cells were cultured in DMEM supplied with 10% (*v/v*) fetal bovine serum and 1% penicillin and streptomycin, and incubated at 37°C under 5% CO_2_. Cells were sub-cultured and used for subsequent experiments when they reached about 80% confluence.

The MTT reduction assay was used to determine the change in gross cell viability. The HT22 cells were usually seeded in 96-well plates at a density of 2,000 cells/well 24 h prior to receiving different experimental treatments. To test the effect of a modulating compound, cells were jointly treated with RSL3 and the compound for 24 h. Afterwards, MTT at a final concentration of 0.5 mg/mL was added to each well, and incubated for 2.5 h at 37°C under 5% CO_2_. After incubation, the MTT-containing medium was removed and 100 μL DMSO was added to each well to dissolve the MTT formazan. Absorbance of the MTT formazan was measured with a UVmax microplate reader (Molecular Device, Palo Alto, CA, USA) at 560 nm.

### siRNA transfection

For siRNA transfection, the procedures were described in our earlier study [12]. Briefly, the siRNAs (at a final concentration of 60 nM) for targeted genes (PDI, iNOS, nNOS, and TrxR) were transfected into HT22 cells 24 h after seeding with Lipofectamine RNA iMAX (Invitrogen). Twenty-four h after siRNA transfection, cells were treated with selected drugs, and subsequently processed for cell viability measurement, fluorescence staining and measurement, and immunoblot analysis.

### Measurement of cellular NO, ROS and lipid-ROS levels

The cellular NO, ROS and lipid-ROS levels were jointly determined using analytical flow cytometry and fluorescence/confocal microscopy. The HT22 cells were seeded in 24-well plates at a density of 5 × 10^4^ cells/well 24 h before treatment. Afterwards, the cells were treated with RSL3 ± different modulating compounds for 6 h or as indicated. It is of note that the 6-h RSL3 treatment time was selected based on our initial analysis of the time course of NO, ROS and lipid-ROS accumulation. Prior to staining with fluorescent probes, cells were washed with HBSS twice. The cellular levels of NO, ROS and lipid-ROS were detected after addition of respective cell-permeant fluorescent probes (10 µM DAF-FM-DA for NO detection, 10 μM DCFH-DA for ROS detection, and 10 μM BODIPY^TM^-581/591-C11 for lipid-ROS detection), which were dissolved in 200 μL phenol red- and serum-free DMEM. The cells were incubated at 37°C for an additional 30 min, and washed twice with HBSS. The fluorescence images of cellular NO and ROS were taken using a fluorescence microscope (AXIO, Carl Zeiss Corporation, Germany). The fluorescence images of the cellular lipid-ROS were captured using a confocal laser scanning microscope (LSM 900; Carl Zeiss, Oberkochen, Germany) and analyzed with the Zen software (Carl Zeiss). The quantitative fluorescence intensity reflecting the cellular accumulation NO, ROS and lipid-ROS was detected using a flow cytometer (Beckman Coulter, Brea, CA, USA). Three or more experiments were usually conducted to confirm the observations, and the average fluorescence intensity (mean ± S.D.) based on the replicate measurements in one representative experiment was calculated with the FlowJo software (FlowJo, LLC, Ashland, USA) and selected for presentation.

For measuring intracellular MDA levels, a MDA Assay Kit was used, and all procedures followed manufacturer’s instructions.

### Measurement of TrxR1 enzyme activity

The TrxR1 enzyme activity was determined using a commercial kit which is based on the measurement of NADPH-dependent reduction of the disulfide bond in the substrate 5,5’-dithiobis-2-nitrobenzoic acid (DTNB). To determine the effect by RSL3 or auranofin on TrxR1’s enzyme activity in cultured HT22 cells, the cells were treated with RSL3 (100 nM) or auranofin (1 μM) for 6 h, and then washed with PBS twice, lysed and centrifuged at 9500 *g* for 10 min (4°C) to collect cellular supernatants. Aliquots of the supernatants were properly diluted and then added into the 96-well plate for determination of TrxR1 enzyme activity according to manufacturer’s instructions. The change in absorbance at 412 nm was recorded using a UVmax microplate reader (Molecular Device, Palo Alto, CA, USA). The relative TrxR1 enzyme activity was calculated as a percentage of the vehicle-treated control cells. To determine the direct inhibition of TrxR1’s catalytic activity by RSL3 and auranofin in the *in-vitro* enzymatic assay, the untreated HT22 cells were washed with PBS twice, lysed and then centrifuged at 9500 *g* for 10 min (4°C) to prepare cellular supernatants which contain the TrxR1 enzyme. Different concentrations of RSL3 and auranofin were included in the incubation, and the enzyme activity in each treatment group was determined by monitoring the change in absorbance at 412 nm once every 2 min for up to 150 min. Here it should be noted that in this enzyme activity assay, it is assumed that the NADPH- dependent specific catalytic activity of TrxR1 is inhibited by high concentrations of auranofin (its *IC*_50_ for TrxR1 is approximately 1.1 nM as determined in this study), and the baseline reading at absorbance 412 nm which cannot be inhibited by 1000 nM auranofin is subtracted.

### Immunoblot analysis

After treatment with RSL3 for different durations as indicated, the HT22 cells were collected by trypsinization and centrifugation, and then lysed in a RIPA buffer containing 1% protease inhibitor cocktail on ice for 30 min. Protein concentrations were determined using the BCA Assay Kit (ThermoFisher). For total nNOS and iNOS analysis (including both monomeric and dimeric forms of nNOS and iNOS), samples were heated at 95°C for 5 min with a reducing buffer before loaded onto the electrophoresis gel. To analyze the monomeric and dimeric forms of nNOS and iNOS, the samples were prepared with a non-reducing buffer and were not heated, and the temperature of the gel was maintained below 15°C during electrophoresis. The proteins were separated using 10% SDS-PAGE gel (for total nNOS and iNOS analysis) or 6% SDS- PAGE gel (for monomeric and dimeric nNOS and iNOS analysis), and then transferred to PVDF membranes. The membranes were incubated for 1 h in 5% skim milk at room temperature, and then incubated with the primary antibody overnight. Afterwards, the membranes were washed three times with TBST at room temperature (10 min each time). The membranes were then incubated with the secondary antibody for 1 h at room temperature and washed with TBST three times before visualization. Desired protein bands were visualized using the chemiluminescence imaging system (Tanon 5200, Shanghai, China).

### Statistical analysis

All quantitative experiments and data described in this study were repeated at least three times. The data were usually presented as mean ± S.D. of multiple replicate measurements from one selected representative experiment. Statistical analysis was conducted using one-way ANOVA, followed by Dunnett’s *post-hoc* tests for multiple comparisons as needed (GraphPad Prism 7, GraphPad Software, La Jolla, CA). Statistical significance was denoted by *P* < 0.05 (* or ^#^) and *P* < 0.01 (** or ^##^) for significant and very significant differences, respectively. In most cases, * and ** denote the comparison for statistical significance between the control group (cells treated with vehicle only) and the cells treated with a chemical (such as RSL3), whereas ^#^ and ^##^ denote the comparison between the cells treated with RSL3 alone and the cells jointly treated with RSL3 and another compound. For Western blot quantification, one representative data is shown.

## RESULTS

### Role of cellular NO, ROS and lipid-ROS accumulation in RSL3-induced ferroptotic cell death

Treatment of HT22 cells with RSL3-induced cell death in time- and dose-dependent manners (**Fig. 1A-1C**). Ferrostatin-1 (Fer-1), a commonly-used lipid-ROS scavenger [13, 14], showed a strong protection against RSL3-induced cytotoxicity in these cells (**Fig. 1D**), which confirms earlier observations [15, 16] reporting that RSL3 induces ferroptotic cell death in cultured HT22 cells.

**Figure 1.**
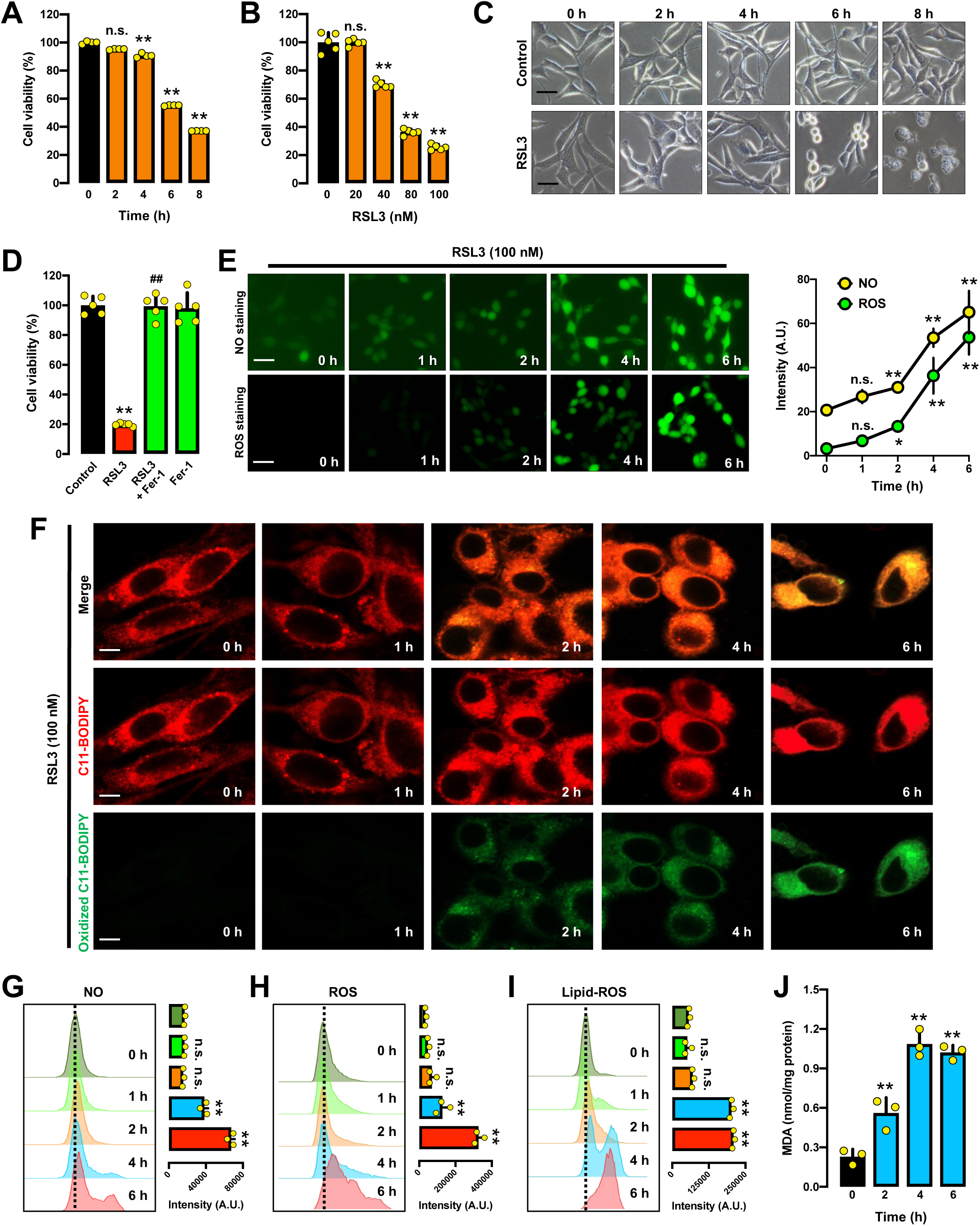
Time- and dose-dependent accumulation of NO, ROS and lipid-ROS in RSL3-induced ferroptosis in HT22 cells. **A, B.** Time- and dose-dependent effect of RSL3 on cell viability (MTT assay, n = 5). The RSL3 concentration used is 100 nM (**A**) and the treatment duration is 24 h (**B**). **C.** Time-dependent change in the gross morphology in RSL3 (100 nM)-treated cells (scale bar = 45 µm). **D.** Protective effect of Fer-1 against RSL3-induced cytotoxicity following treatment of cells with RSL3 (100 nM) ± Fer-1 (1 µM) for 24 h (MTT assay, n = 5). **E.** Time-dependent accumulation of cellular NO and ROS (fluorescence microscopy images, scale bar = 60 µm) following RSL3 treatment, and the quantitative intensity values are shown in the right panel (n = 3). **F.** Time-dependent accumulation of cellular lipid-ROS (confocal microscopy images) in RSL3-treated cells (scale bar = 10 µm). **G, H, I.** Time-dependent accumulation of cellular NO (**G**), ROS (**H**) and lipid-ROS (**I**) in RSL3-treated cells (analytical flow cytometry). The quantitative intensity values are shown in the right panel (n = 3). **J.** Time-dependent change in cellular MDA levels in RSL3-treated cells (n = 3). Each value represents the mean ± S.D. (* *P* < 0.05; ** or ^##^ *P* < 0.01; n.s. not significant).

Our recent studies showed that during erastin-induced, GSH depletion-associated ferroptotic cell death in HT22 cells, there was a time-dependent sequential induction of NO and ROS/lipid-ROS accumulation [12]. To probe whether RSL3-induced ferroptosis in HT22 cells is also related to cellular NO and ROS/lipid-ROS accumulation, we determined the time-dependent change in cellular NO, ROS and lipid-ROS levels following RSL3 treatment (**Fig. 1E-1J**). Based on measurements using fluorescence microscopy and flow cytometry, we found that treatment of HT22 cells with RSL3 elicited a time-dependent increase in cellular NO level (**Fig. 1E, 1G, Fig. S1A**). Similarly, the cellular levels of ROS and lipid-ROS were also increased in a time- and dose- dependent manner in RSL3-treated cells (**Fig. 1E, 1H, Fig. S1B** for ROS; **Fig. 1F, 1I** for lipid-ROS). Malondialdehyde (MDA), one of the end products of lipid peroxidation and often used as an indicator of cellular lipid peroxidation [17], was increased by RSL3 treatment in a time-dependent manner (**Fig. 1J**). It was observed that NO accumulation occurs at a very similar time as does ROS/lipid-ROS accumulation.

To determine whether NO accumulation subsequently leads to ROS and lipid-ROS accumulation and ultimately ferroptotic cell death, we studied the effect of 2-(4-carboxyphenyl)-4,4,5,5-tetramethylimidazoline-1-oxyl-3-oxide (cPTIO), a NO scavenger, in RSL3-treated cells [18]. cPTIO effectively suppressed RSL3-induced accumulation of NO (**Fig. 2A**), ROS (**Fig. 2B**) and lipid-ROS (**Fig. 2C**) in HT22 cells. In addition, cPTIO also attenuated RSL3-induced cell death (**Fig. 2D**).

**Figure 2.**
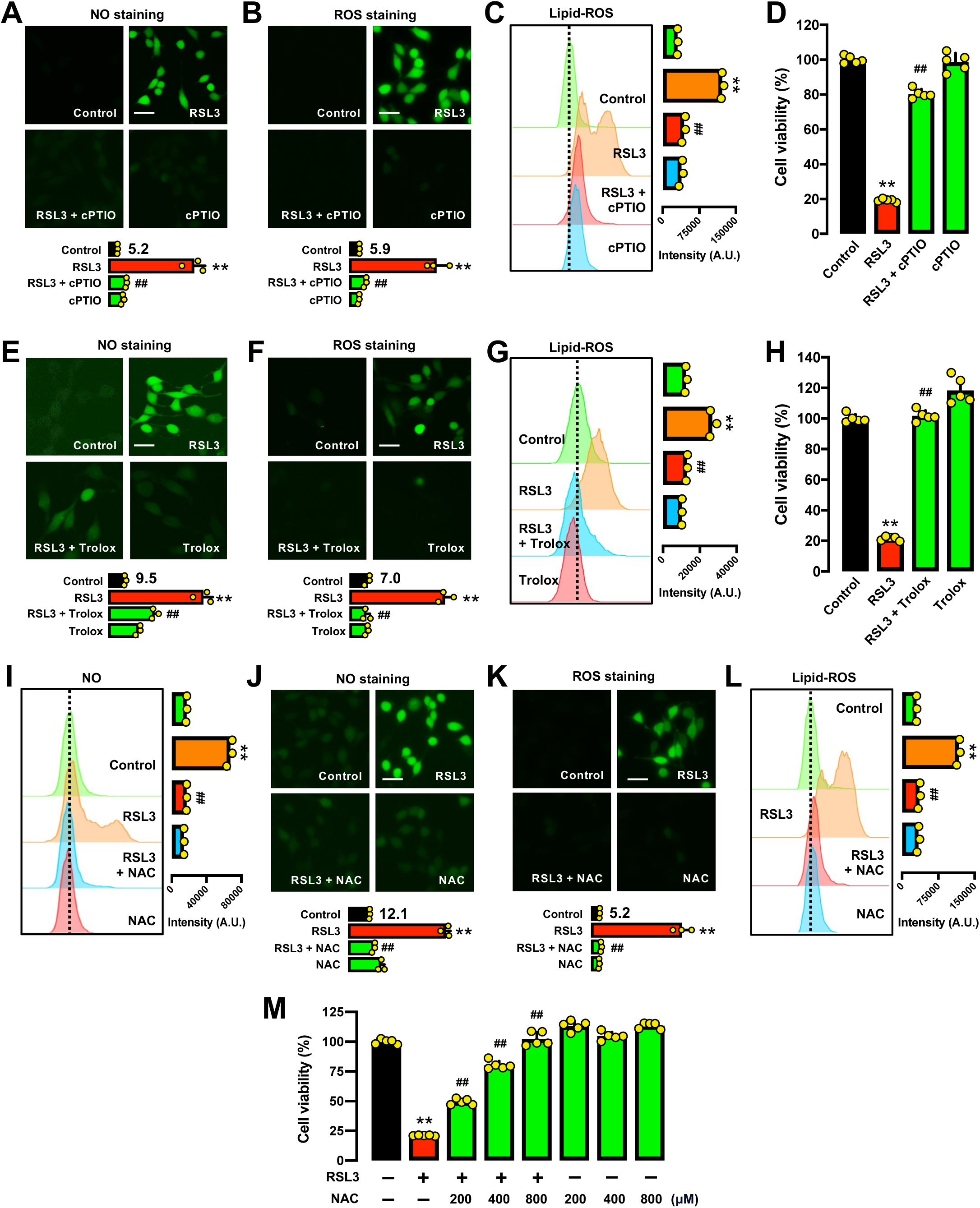
Effect of free radical scavengers on RSL3-induced cytotoxicity and cellular accumulation of NO, ROS and lipid-ROS in HT22 cells. **A, B, C.** Abrogation by cPTIO of RSL3-induced accumulation of cellular NO (**A**), ROS (**B**) and lipid-ROS (**C**). Cells were treated with RSL3 (100 nM) ± cPTIO (200 µM) for 6 h, and then stained for NO, ROS and lipid-ROS (**A, B** for fluorescence microscopy; scale bar = 60 µm; **C** for analytical flow cytometry). The quantitative intensity values are shown in the lower panel (n = 3; the mean value for the control group is labeled next to the bar). **D.** Protective effect of cPTIO against RSL3-induced cytotoxicity following treatment of cells with RSL3 (100 nM) ± cPTIO (200 µM) for 24 h (MTT assay, n = 5). **E, F, G.** Abrogation by Trolox of RSL3-induced accumulation of cellular NO (**E**), ROS (**F**) and lipid-ROS (**G**). Cells were treated with RSL3 (100 nM) ± Trolox (100 µM) for 6 h, and then stained for NO, ROS and lipid-ROS (**E, F** for fluorescence microscopy; scale bar = 60 µm; **G** for analytical flow cytometry). The quantitative intensity values are shown in the lower panel (n = 3; the mean value for the control group is labeled next to the bar). **H.** Protective effect of Trolox against RSL3-induced cytotoxicity following treatment of cells with RSL3 (100 nM) ± Trolox (100 µM) for 24 h (MTT assay, n = 5). **I, J, K, L.** Abrogation by NAC of RSL3-induced accumulation of cellular NO (**I, J**), ROS (**K**) and lipid-ROS (**L**). Cells were treated with RSL3 (100 nM) ± NAC (800 µM) for 6 h, and then stained for NO, ROS and lipid-ROS (**J, K** for fluorescence microscopy; scale bar = 60 µm; **I, L** for analytical flow cytometry). The quantitative intensity values are shown in the lower panel (n = 3; the mean value for the control group is labeled next to the bar). **M.** Protective effect of NAC against RSL3-induced cytotoxicity following treatment of cells with RSL3 (100 nM) ± NAC (200, 400 and 800 µM) for 24 h (MTT assay, n = 5). Each value represents the mean ± S.D. (** or ^##^ *P* < 0.01).

6-Hydroxy-2,5,7,8-tetramethylchroman-2-carboxylic acid (Trolox), which has a strong ROS-scavenging activity plus a weaker NO-scavenging activity [19], partially abrogated RSL3-induced accumulation of NO (**Fig. 2E**) but strongly abrogated the accumulation of ROS (**Fig. 2F**) and lipid-ROS (**Fig. 2G**), and these effects were associated with a very strong protection against RSL3-induced ferroptotic cell death (**Fig. 2H**).

NAC (*N*-acetyl-*L* -cysteine), another commonly-used antioxidant [20], very strongly abrogated RSL3-induced accumulation of cellular NO (**Fig. 2I, 2J**), ROS (**Fig. 2K**) and lipid-ROS (**Fig. 2L**), and these effects were also associated with a strong protection against RSL3-induced cell death (**Fig. 2M**).

### Enhancement of RSL3-induced ferroptotic cell death by sodium nitroprusside

Sodium nitroprusside (SNP) is a NO donor and can rapidly release NO inside the cells [21]. To provide further support for an upstream role of NO in RSL3-induced ROS/lipid-ROS accumulation and ferroptosis, we sought to determine the effect of SNP on RSL3-induced ROS/lipid-ROS accumulation and ferroptosis in HT22 cells. First, treatment of HT22 cells with SNP alone caused a rapid rise in cellular NO level (**Fig. 3A, 3B, 3C**), which was accompanied by increase in cellular ROS level (**Fig 3A, 3B, 3D**) and lipid-ROS level (**Fig. 3E**). It is of note while NO accumulation peaked within 0.5 h after SNP exposure, the accumulation of ROS and lipid-ROS lagged slightly behind NO increase, reaching the peak at around 1 h after SNP exposure. In addition, treatment of HT22 cells with SNP alone also elicited a dose-dependent cytotoxicity (**Fig. 3F**).

**Figure 3.**
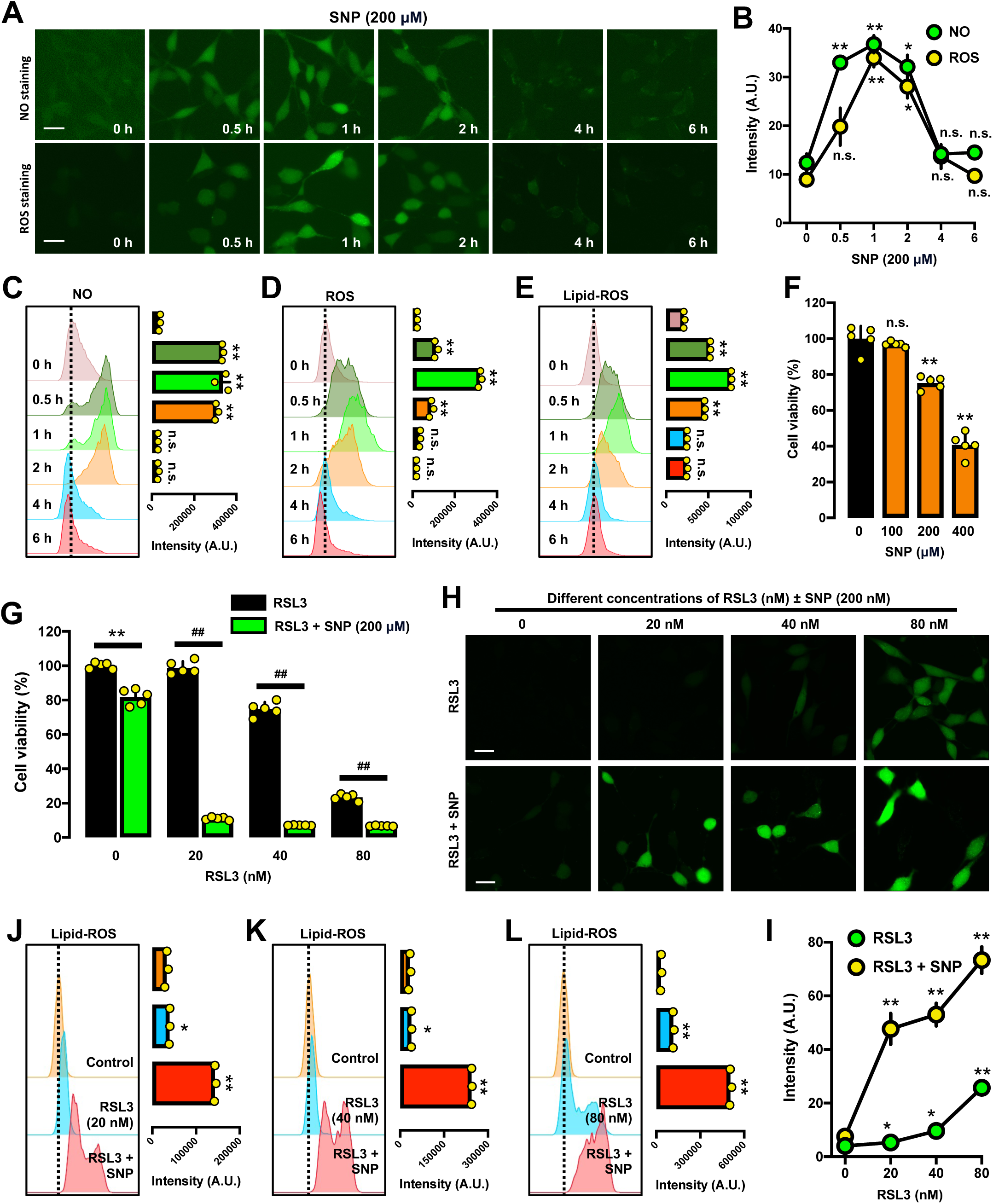
Effect of SNP on RSL3-induced NO, ROS and lipid-ROS accumulation in HT22 cells. **A, B.** Levels of NO and ROS after treatment with SNP (200 μM) for different durations (0.5, 1, 2, 4 and 6 h; scale bar = 50 µm) (**A**). The quantitative intensity values are shown in the right panel (n = 3) (**B**) **C, D, E.** Time-dependent accumulation of cellular NO (**C**), ROS (**D**) and lipid-ROS (**E**) following SNP treatment (analytical flow cytometry) and the quantitative intensity values are in the right panel (n = 3). **F.** Dose-dependent effect of SNP on cell viability (MTT assay, n = 5). **G.** Cell viability change following treatment with SNP (200 μM) and different concentrations of RSL3 for 24 h (MTT assay, n = 5). **H, I, J, K, L.** Accumulation of ROS (**H, I**) and lipid-ROS (**J, K, L**) after treatment with RSL3 (100 nM) ± SNP (200 μM) (**H** for fluorescence microscopy; scale bar = 60 µm; **J, K, L** for analytical flow cytometry). The quantitative intensity values for **H** are shown in the lower panel (**I**) (n = 3). Each value represents the mean ± S.D. (* or ^#^ *P* < 0.05; ** *P* < 0.01; n.s., not significant).

It is of note that joint treatment of HT22 cells with RSL3 + SNP (200 μM) drastically sensitized HT22 cells to RSL3-induced ferroptotic cell death (**Fig. 3G**). The observed enhancement of RSL3’s cytotoxicity by SNP was associated in a parallel manner with a drastically-enhanced accumulation of cellular ROS (**Fig. 3H, 3I**) and lipid-ROS (**Fig. 3J, 3K, 3L**) in cells jointly treated with RSL3 + SNP. Together, these results provide additional support for the hypothesis that RSL3-induced NO accumulation is an important determinant for accumulation of cellular ROS and lipid-ROS, and subsequently ferroptotic cell death.

### Role of NOS induction and activation in RSL3-induced ferroptotic cell death

It is known that the generation of cellular NO is mostly catalyzed by the nitric oxide synthase (NOS), which uses *L* -arginine as its primary substrate [22–24]. There are three known subtypes of NOS, namely, nNOS, iNOS and eNOS [22–24]. Our earlier study showed that while nNOS was readily detected in untreated HT22 cells, iNOS was mostly undetectable [25]. In this study, we found that treatment of HT22 cells with increasing concentrations of RSL3 for different durations did not significantly affect total cellular nNOS protein level (including both monomer and dimer forms) (**Fig. 4A**). By contrast, while little or no iNOS band was detected in vehicle-treated control HT22 cells as reported earlier [25], treatment of these cells with RSL3 significantly increased the total cellular iNOS level in a time- and dose-dependent manner (**Fig. 4A**).

**Figure 4.**
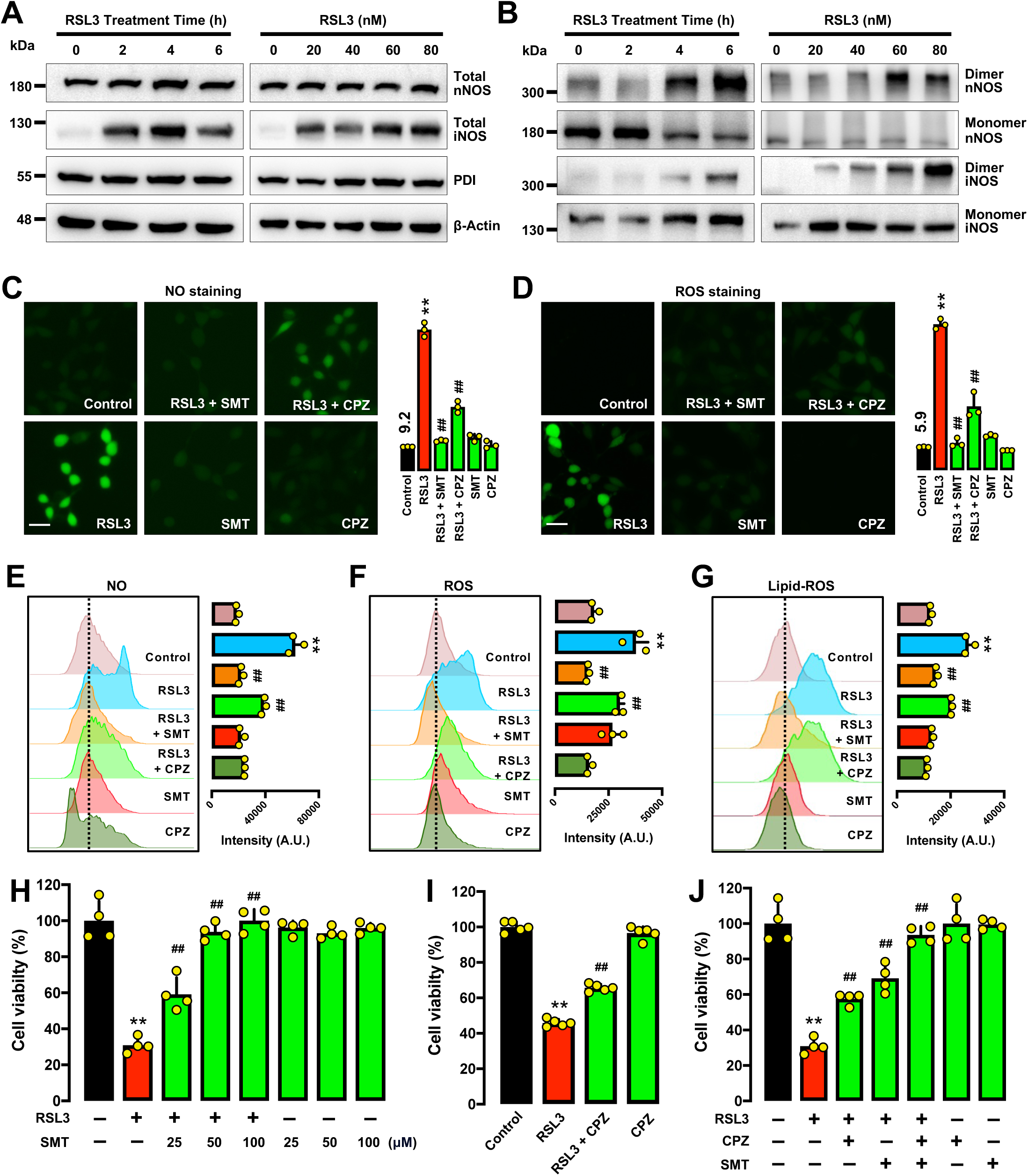
RSL3-induced NOS activation in HT22 cells. **A.** Levels of total iNOS and nNOS proteins following treatment of cells with RSL3 (100 nM) for the indicated time intervals, or with increasing concentrations of RSL3 for 6 h. **B.** Levels of the dimer and monomer iNOS and nNOS proteins following treatment of cells with RSL3 (100 nM) for the indicated time intervals, or with increasing concentrations of RSL3 for 6 h. **C, D, E, F, G.** Abrogation by SMT and CPZ of RSL3-induced accumulation of cellular NO (**C, E**), ROS (**D, F**) and lipid-ROS (**G**). Cells were treated with RSL3 (100 nM) ± SMT (50 µM) or RSL3 (100 nM) ± CPZ (40 µM) for 6 h, and then stained for NO, ROS and lipid- ROS (**C, D** for fluorescence microscopy; scale bar = 60 µm; **E, F, G** for analytical flow cytometry). The quantitative intensity values are shown on the right panel (n = 3; the mean value for the control group is labeled next to the bar). **H.** Protective effect of SMT against RSL3-induced cytotoxicity following treatment of cells with RSL3 (100 nM) ± SMT (25, 50, 100 µM) for 24 h (MTT assay, n = 4). **I.** Protective effect of SMT against RSL3-induced cytotoxicity following treatment of cells with RSL3 (100 nM) ± CPZ (40 µM) for 24 h (MTT assay, n = 5). **J.** Protective effect of SMT + CPZ against RSL3-induced cytotoxicity following treatment of cells with RSL3 (100 nM) ± SMT (25 µM) and CPZ (12.5 µM) for 24 h (MTT assay, n = 4). Each value represents the mean ± S.D. (** or ^##^ *P* < 0.01).

The catalytic activity of NOS is activated through the formation of NOS dimers [22–24], which are stabilized by the formation of an intermolecular disulfide bond [26]. Our earlier studies showed that the NOS proteins are activated to form dimers in erastin-treated cells [12, 27]. Next, we sought to determine whether nNOS and iNOS were activated (*i.e.*, dimerized) in RSL3-treated HT22 cells. For nNOS, treatment of HT22 cells with RSL3 significantly increased the formation of nNOS dimers (**Fig. 4B**), but the total nNOS protein level was not significantly changed (**Fig. 4A**). For iNOS, RSL3 not only increased iNOS activation (dimerization) in a time- and dose-dependent manner (**Fig. 4B**), but it also markedly increased the total iNOS protein level (**Fig. 4A**).

Next, we determined the modulating effect of S-methylisothiourea (SMT), a preferential inhibitor of iNOS [28, 29], and chlorpromazine (CPZ), a preferential inhibitor of nNOS [30], on RSL3-induced cytotoxicity in HT22 cells. We found that joint treatment of HT22 cells with SMT abrogated RSL3-induced accumulation of cellular NO (**Fig. 4C, 4E**), ROS (**Fig. 4D, 4F**) and lipid-ROS (**Fig. 4G**), and these effects were associated with a strong protection (∼100% restoration of cell viability) against RSL3-induced ferroptotic cell death (**Fig. 4H**). In comparison, the effect of CPZ against RSL3- induced accumulation of NO (**Fig. 4C, 4E**), ROS (**Fig. 4D, 4F**) and lipid-ROS (**Fig. 4G**) was markedly weaker, which is also associated with a much weaker protection against RSL3-induced cell death (**Fig. 4I**). The combined use of CPZ and SMT at lower concentrations which only produces a partial cytoprotection when each chemical was used alone exerted a stronger protective effect (**Fig. 4J**).

Lastly, we also evaluated in this study the contributing role of nNOS and iNOS in mediating RSL3-induced ferroptotic cell death by using the siRNA approach to selectively knock down their expression in HT22 cells. The efficiency of iNOS and nNOS knockdown was confirmed by measuring the change in cellular nNOS and iNOS protein levels (**Fig. 5A**). Knockdown of iNOS abrogated RSL3-induced increase in iNOS protein levels (**Fig. 5A**), and it also exerted a significant protection against RSL3-induced cytotoxicity (**Fig. 5B**). By contrast, selective knockdown of nNOS in HT22 cells did not show a significant protective effect (**Fig. 5C**). It is of note that the nNOS knockdown experiment was repeated multiple times, and similar results were obtained which showed a lack of significant protective effect against RSL3-induced cytotoxicity.

**Figure 5.**
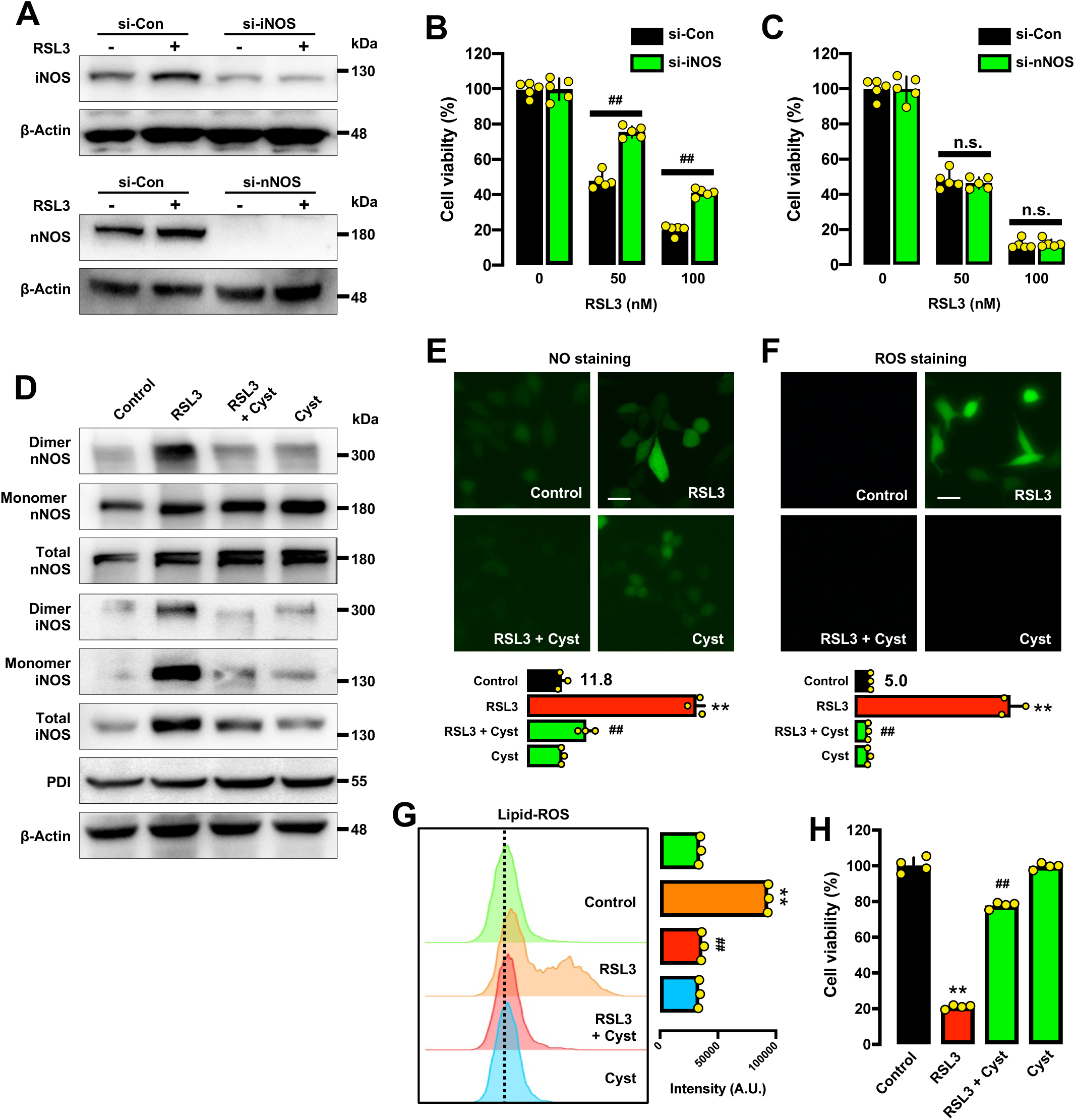
Protective effect of cystamine (a PDI inhibitor) against RSL3-induced ferroptosis in HT22 cells. **A.** Effectiveness of iNOS-siRNAs and nNOS-siRNAs in reducing cellular iNOS and nNOS protein levels, respectively. Cells were transfected with iNOS-siRNAs or nNOS-siRNAs for 48 h prior to treatment of cells with RSL3 (50 and 100 nM) for an additional 6 h **B, C.** Effect of iNOS (**B**) and nNOS (**C**) knockdown on RSL3-induced cytotoxicity. Cells were transfected with iNOS-siRNAs or nNOS-siRNAs for 24 h prior to treatment of cells with RSL3 (100 nM) for an additional 24 h. **D.** Levels of the dimer and monomer iNOS and nNOS proteins, and the levels of the total iNOS and nNOS proteins after treatment of cells with RSL3 (100 nM) ± cystamine (100 nM) for 6 h. **E, F, G.** Abrogation by cystamine of RSL3-induced accumulation of cellular NO (**E**), ROS (**F**) and lipid-ROS (**G**). Cells were treated with RSL3 (100 nM) ± cystamine (100 µM) for 6 h, and then stained for NO, ROS and lipid-ROS (**E, F** for fluorescence microscopy; scale bar = 60 µm; **G** for analytical flow cytometry). The quantitative intensity values are shown in the lower panel (n = 3; the mean value for the control group is labeled next to the bar). **H.** Protective effect of cystamine against RSL3-induced cytotoxicity following treatment of cells with RSL3 (100 nM) ± cystamine (100 µM) for 24 h (MTT assay, n = 4). Each value represents the mean ± S.D. (** or ^##^ *P* < 0.01; n.s. not significant).

Together, these results indicate that NO accumulation resulting from the induction and activation (dimerization) of iNOS and nNOS in RSL3-treated cells is involved in mediating RSL3-induced ferroptosis in HT22 cells. It is apparent that iNOS plays a more dominant role than nNOS in this process.

### PDI mediates NOS dimerization in RSL3-treated cells

In this study, we also sought to determine whether PDI can catalyze nNOS and iNOS dimerization in RSL3-treated HT22 cells. First, we found that the cellular level of total PDI protein was not significantly affected by RSL3 treatment (**Fig. 4A**). To provide support for the hypothesis that PDI is responsible for catalyzing nNOS and iNOS dimerization, we determined the effect of cystamine on RSL3-induced ferroptosis. Cystamine is an organic disulfide molecule capable of inhibiting PDI’s enzymatic activity through covalent modification of the cysteine residue(s) in its catalytic site [31, 32]. For nNOS, it was observed that treatment of HT22 cells with RSL3 stimulated its activation (dimerization), and joint treatment with cystamine abrogated RSL3-induced nNOS dimerization (**Fig. 5D**). For iNOS, RSL3 markedly increased iNOS total protein level and its dimer formation in HT22 cells (**Fig. 5D**), and joint treatment with cystamine abolished RSL3-induced increase in total iNOS protein level and dimerization (**Fig. 5D**). These effects of cystamine on nNOS and iNOS dimerization and on total iNOS protein level in RSL3-treated cells were associated with a strong abrogation of NO (**Fig. 5E**), ROS (**Fig. 5F**) and lipid-ROS (**Fig. 5G**) accumulation, as well as a strong protection against RSL3-induced ferroptotic cell death (**Fig. 5H**).

We recently have shown that 4-hydroxyestrone (4-OH-E_1_) is an endogenous estrogen metabolite that can effectively inhibit the enzymatic activity of PDI [33]. We found that joint treatment of HT22 cells with 4-OH-E_1_ abrogated RSL3-induced increase in total iNOS protein levels (**Fig. 6A**). Because the nNOS dimers are relatively easier to detect than the iNOS dimers, we used the nNOS dimers as a representative parameter to reflect PDI-mediated NOS dimerization (activation) in RSL3-treated HT22 cells. We found that joint treatment of HT22 cells with 4-OH-E_1_ abrogated RSL3-induced nNOS dimerization (**Fig. 6A**). This effect was associated with decreased accumulation of cellular NO (**Fig. 6B**), ROS (**Fig. 6C**) and lipid-ROS (**Fig. 6D**), together with increased cell survival (**Fig. 6E**).

**Figure 6.**
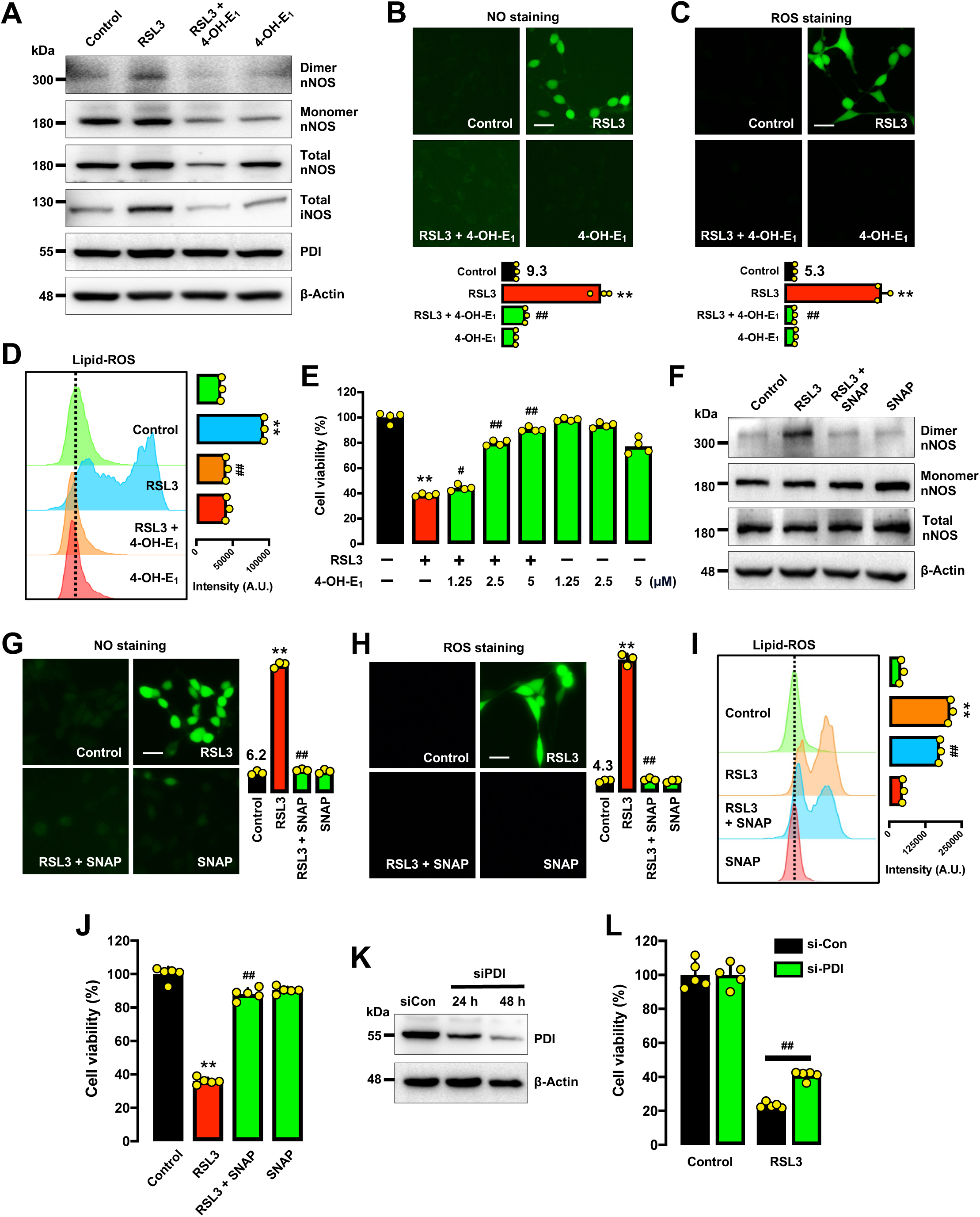
4-OH-E_1_ and SNAP (a S-nitrosylating agent) protect against RSL3-induced ferroptosis in HT22 cells. **A.** Levels of the dimer and monomer nNOS proteins, and the levels of total iNOS and nNOS proteins after treatment of cells with RSL3 (100 nM) ± 4-OH-E_1_ (5 μM) for 6 h. **B, C, D.** Abrogation by 4-OH-E_1_ of RSL3-induced accumulation of cellular NO (**B**), ROS (**C**) and lipid-ROS (**D**). Cells were treated with RSL3 (100 nM) ± 4-OH-E_1_ (5 µM) for 6 h, and then stained for NO, ROS and lipid-ROS (**B, C** for fluorescence microscopy; scale bar = 65 µm; **D** for analytical flow cytometry). The quantitative intensity values are shown in the lower panel (n = 3; the mean value for the control group is labeled next to the bar). **E.** Protective effect of 4-OH-E_1_ against RSL3-induced cytotoxicity following treatment of cells with RSL3 (100 nM) ± 4-OH-E_1_ (1.25, 2.5 and 5 µM) for 24 h (MTT assay, n = 4). **F.** Levels of the dimer and monomer nNOS proteins and the total nNOS protein after treatment of cells with RSL3 (100 nM) ± SNAP (200 μM) for 6 h. **G, H, I.** Abrogation by SNAP of RSL3-induced accumulation of cellular NO (**G**), ROS (**H**) and lipid-ROS (**I**). Cells were treated with RSL3 (100 nM) ± SNAP (200 µM) for 6 h, and then stained for NO, ROS and lipid-ROS (**G, H** for fluorescence microscopy; scale bar = 65 µm; **I** for analytical flow cytometry). The quantitative intensity values are shown in the right panel (n = 3; the mean value for the control group is labeled next to the bar). **J**. Protective effect of SNAP against RSL3-induced cytotoxicity following treatment of cells with RSL3 (100 nM) ± SNAP (200 µM) for 24 h (MTT assay, n = 5). **K.** Effectiveness of PDI-siRNAs in reducing cellular PDI protein level. Cells were transfected with PDI-siRNAs for 24 and 48 h, respectively, and the PDI protein levels were determined by Western blotting. **L.** Protective effect of PDI knockdown on RSL3-induced cytotoxicity. Cells were transfected with PDI-siRNAs for 24 h prior to treatment of cells with RSL3 (100 nM) for an additional 24 h. Each value represents the mean ± S.D. (^#^ *P* < 0.05; ** or ^##^ *P* < 0.01).

Our earlier study showed that PDI in untreated HT22 cells is mostly present in the catalytically-inactive *S*-nitrosylated form, and it undergoes *S*-denitrosylation following GSH depletion [25]. To probe the involvement of PDI *S*-denitrosylation in mediating RSL3-induced cytotoxicity in HT22 cells, we studied the effect of *S*-nitroso-*N*- acetylpenicillamine (SNAP), a protein *S*-nitrosylating agent [34]. We found that joint treatment of HT22 cells with SNAP abrogated RSL3-induced NOS activation (based on detection of nNOS dimer; **Fig. 6F**), and this effect was associated with reduced accumulation of NO (**Fig. 6G**), ROS (**Fig. 6H**), lipid-ROS (**Fig. 6I**), and protection against ferroptotic cell death (**Fig. 6J**). These results indicate that *S*-nitrosylation of PDI is indeed associated with suppression of PDI activity, reduction in NOS dimerization, reduction in cellular NO, ROS and lipid-ROS levels, and ultimately, protection from ferroptosis.

Based on the consistent observations with different PDI inhibitors, next we also tried to probe PDI’s effect by using specific siRNAs to selectively knockdown its expression. The effectiveness of PDI knockdown was assessed by Western blotting of its cellular protein levels (**Fig. 6K**). As expected, PDI knockdown elicited a partial protection against RSL3-induced cytotoxicity (**Fig. 6L**), but the degree of protection in PDI-knockdown cells is smaller than expected, which likely is due to the fact that a transient PDI knockdown may not last long enough for the cell viability experiment.

Collectively, these results still clearly show that PDI is a key upstream mediator of RSL3-induced ferroptosis by catalyzing iNOS dimerization in RSL3-treated HT22 cells, which then leads to accumulation of cellular NO and ROS/lipid-ROS and ultimately, oxidative cell death.

### Role of TrxR1 in RSL3-induced ferroptosis

A recent study reported that RSL3 can inhibit the catalytic activity of TrxR1 [5]. Since PDI belongs to the thioredoxin superfamily and a substrate of TrxR1 [35, 36], next we sought to determine the potential role of TrxR1 in RSL3-induced ferroptotic cell death. First, we used the TrxR1-siRNAs to transiently knock down its expression in HT22 cells. The effectiveness of TrxR1 knockdown was confirmed by Western blot analysis of TrxR1 protein levels (**Fig. 7A**). While TrxR1 knockdown alone (in the absence of RSL3) reduced the baseline cell viability compared to cells transfected with the control siRNAs alone (**Fig. S2A**), TrxR1 knockdown markedly reduced the sensitivity of cells to the cytotoxicity induced by RSL3 and auranofin (a prototypical inhibitor of TrxR1 [5, 37, 38]) (**Fig. 7B**).

**Figure 7.**
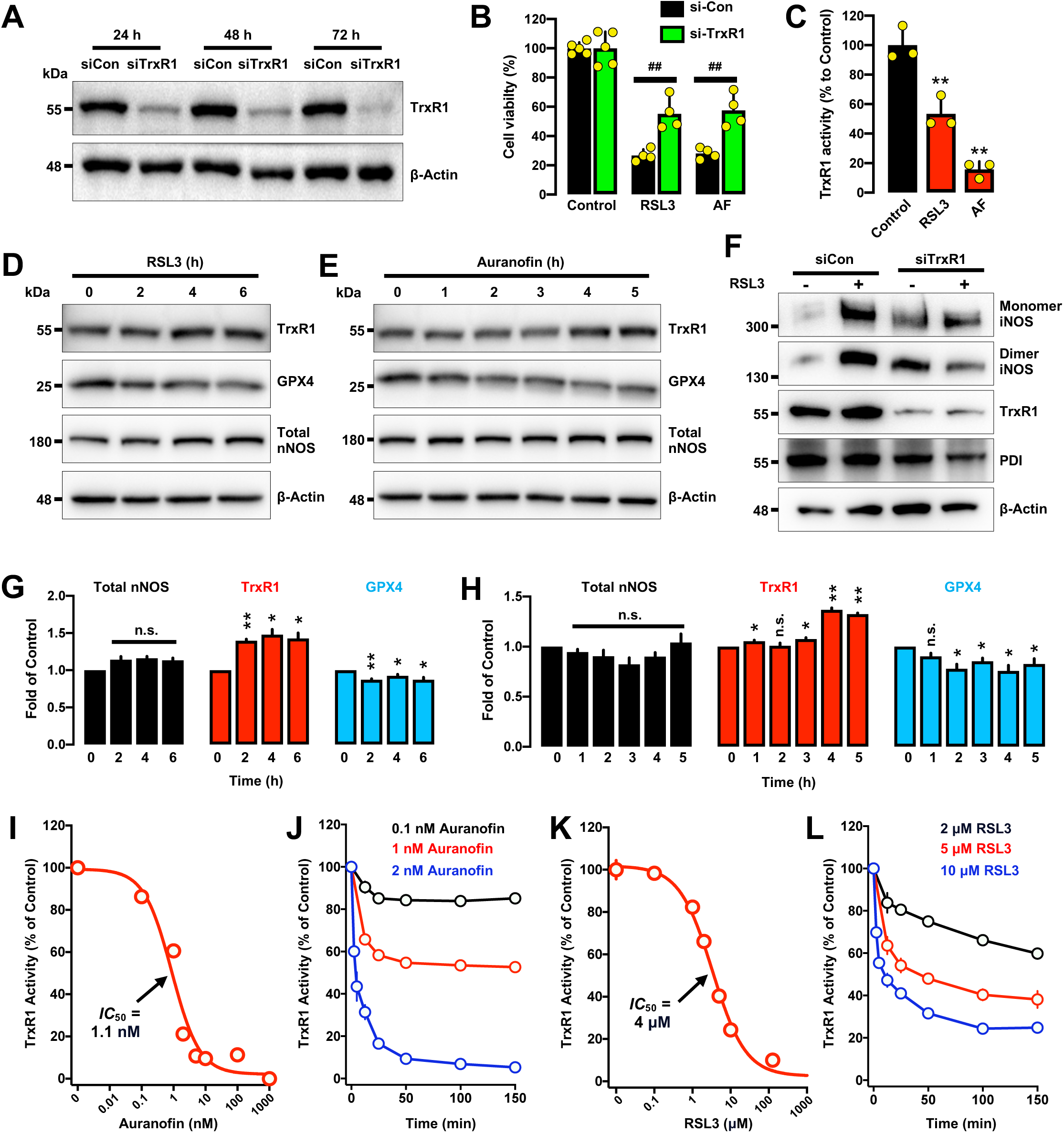
Role of TrxR1 in RSL3-induced ferroptosis in HT22 cells. **A.** Effectiveness of TrxR1-siRNAs in reducing cellular TrxR1 protein level. Cells were transfected with TrxR1-siRNAs for 24, 48 and 72 h, and the TrxR1 protein levels were determined by Western blotting. **B.** Effect of TrxR1 knockdown on RSL3- and auranofin-induced cytotoxicity. Cells were transfected with TrxR1-siRNAs for 24 h prior to the treatment of RSL3 (100 nM) or auranofin (1 µM) for an additional 24 h, and then cell viability was determined (MTT assay, n = 4). **C.** Levels of TrxR1 enzyme activity in cells following treatment with RSL3 (100 nM) or auranofin (1 μM) for 6 h. Cellular supernatants were prepared from different treatment groups (using same numbers of cells), and then the TrxR1 enzyme activity was assayed. The relative TrxR1 activity in the RSL3 and auranofin groups was presented as a percentage of the relative TrxR1 activity in the vehicle-treated control group. **D, E.** Levels of TrxR1, GPX4 and total nNOS proteins following treatment with 100 nM RSL3 (**D**) or 1 μM auranofin (**E**) for different durations as indicated. **F.** Levels of iNOS monomer, iNOS dimer, TrxR1 and PDI proteins in RSL3-treated cells with TrxR1 knockdown. Cells were transfected with control-siRNAs or TrxR1 siRNAs for 24 h and then treated with 100 nM RSL3 for an additional 6 h. **G, H.** Relative protein band intensity of TrxR1, GPX4 and total nNOS proteins following treatment with 100 nM RSL3 (**G**) or 1 μM auranofin (**H**) for different durations as indicated. The protein bands were determined by densitometry counting (the ImageJ software). The relative intensity of a given protein was normalized to the corresponding β-actin intensity, and the relative protein level in a given treatment group is presented relative to the vehicle-treated control group. **H, I, J, K.** Concentration- and incubation time-dependent inhibition of TrxR1’s enzyme activity *in vitro* by auranofin (**H, I**) and RSL3 (**J, K**). The TrxR1 enzyme activity in the presence of an inhibitor (auranofin or RSL3) was presented as % of the control activity in the absence of an inhibitor. Each value is the mean ± S.D. (* *P* < 0.05; ** or ^##^ *P* < 0.01; n.s., not significant).

We also determined whether the TrxR1 activity in HT22 cells was inhibited by treatment of cells with RSL3 or auranofin. The HT22 cells were first treated with 0.1 μM RSL3 or 1 μM auranofin for 5 h, then the cells were washed and harvested for preparation of cell lysates. Even after all these preparative steps, the TrxR1 activity contained in the cell lysates was still markedly reduced in RSL3- or auranofin-treated cells (**Fig. 7C**); in comparison, the TrxR1 protein levels remained largely unchanged or slightly increased in these cells (**Fig. 7D, 7E, 7G, 7H**). Jointly, these observations suggest that RSL3 and auranofin can bind to TrxR1 very tightly (most likely covalently) to exert a strong and sustained inhibition of the enzyme activity.

Next, we determined the iNOS dimer level in RSL3-treated HT22 cells with prior TrxR1 knockdown. As shown in **Fig. 7F**, TrxR1 knockdown alone (without RSL3 treatment) caused a significant increase in iNOS dimer level, but it significantly abrogated RSL3-induced increase in iNOS dimer formation. Notably, the cellular PDI level appeared to be also reduced in TrxR1-knockdown cells when it was treated with RSL3 (**Fig. 7F**). These results suggest that TrxR1 plays an important role in meditating PDI-catalyzed iNOS dimerization and subsequent ferroptotic cell death in HT22 cells.

It is of note that in this study, we also probed the effect of jointly knocking down PDI and TrxR1 in HT22 cells and then determined its effect on RSL3-induced cell death. We found that joint knockdown of PDI and TrxR1 reduced the sensitivity to RSL3 cytotoxicity in a similar manner as what was seen with PDI knockdown alone (**Fig. S2B**). This result is expected because PDI is a substrate of TrxR1, and when PDI is mostly absent (resulting from its knockdown), there is not much effect left to be seen even by further knocking down TrxR1.

Next, we determined the ability of RSL3 and auranofin to directly inhibit the enzymatic activity of TrxR1 in biochemical assays. The whole cell lysates were prepared from untreated HT22 cells, and then the lysates (which contain TrxR1 enzyme) were used to assay the TrxR1 enzyme activity in the presence or absence of RSL3 or auranofin. We found that the TrxR1 enzyme activity could be very potently inhibited in a concentration-dependent manner by the presence of auranofin, with an apparent *IC*_50_ of approximately 1.1 nM (**Fig. 7I**). It is of note that the TrxR1 enzyme inhibition by auranofin became stronger but with prolonged incubation time until it reached a plateau (**Fig. 7J**). This pattern of inhibition is consistent with the earlier observation that auranofin can covalently inhibit TrxR1 [39]. Similarly, the TrxR1 enzyme activity was also dose-dependently inhibited by RSL3, with an apparent *IC*_50_ of approximately 4 μM (**Fig. 7K**), which is in agreement with the results from a recent study when the purified TrxR1 enzyme was used in the *in vitro* assay [5]. It is of note that the overall pattern of TrxR1 inhibition by RSL3 is similar to the inhibition by auranofin, *i.e.*, a stronger inhibition was observed with prolonged reaction time, until it reached a plateau (**Fig. 7L**).

Next, we determined whether auranofin can induce cytotoxicity in HT22 cells. Treatment of HT22 cells with auranofin alone elicited a concentration-dependent cytotoxicity (**Fig. 8A**). The *IC* _50_ of auranofin’s cytotoxicity is 0.8 μM, which is much higher than its *IC*_50_ for TrxR1 inhibition. In addition, auranofin-induced cell death was not protected by Fer-1 (**Fig. 8B**) or the iron chelator DFO (**Fig. 8**), two commonly-used protective agents for chemically-induced ferroptosis. Similarly, while auranofin (at the 1 μM concentration) elicited accumulation of cellular NO (**Fig. 8D**) and ROS (**Fig. 8E**), it did not elicit a significant accumulation of lipid-ROS (**Fig. 8F**). Together, these observations suggest that the cell death induced by auranofin at high concentrations (much higher than its *IC*_50_ for TrxR1 inhibition) is not ferroptosis. In addition, the apoptosis inhibitor z-VAD-FMK and necrosis inhibitor Nec-1 also failed to prevent auranofin-induced cell death (**Fig. 8G, 8H**), indicating that auranofin-induced cytotoxicity in HT22 cells is also not typical apoptosis or necrosis.

**Figure 8.**
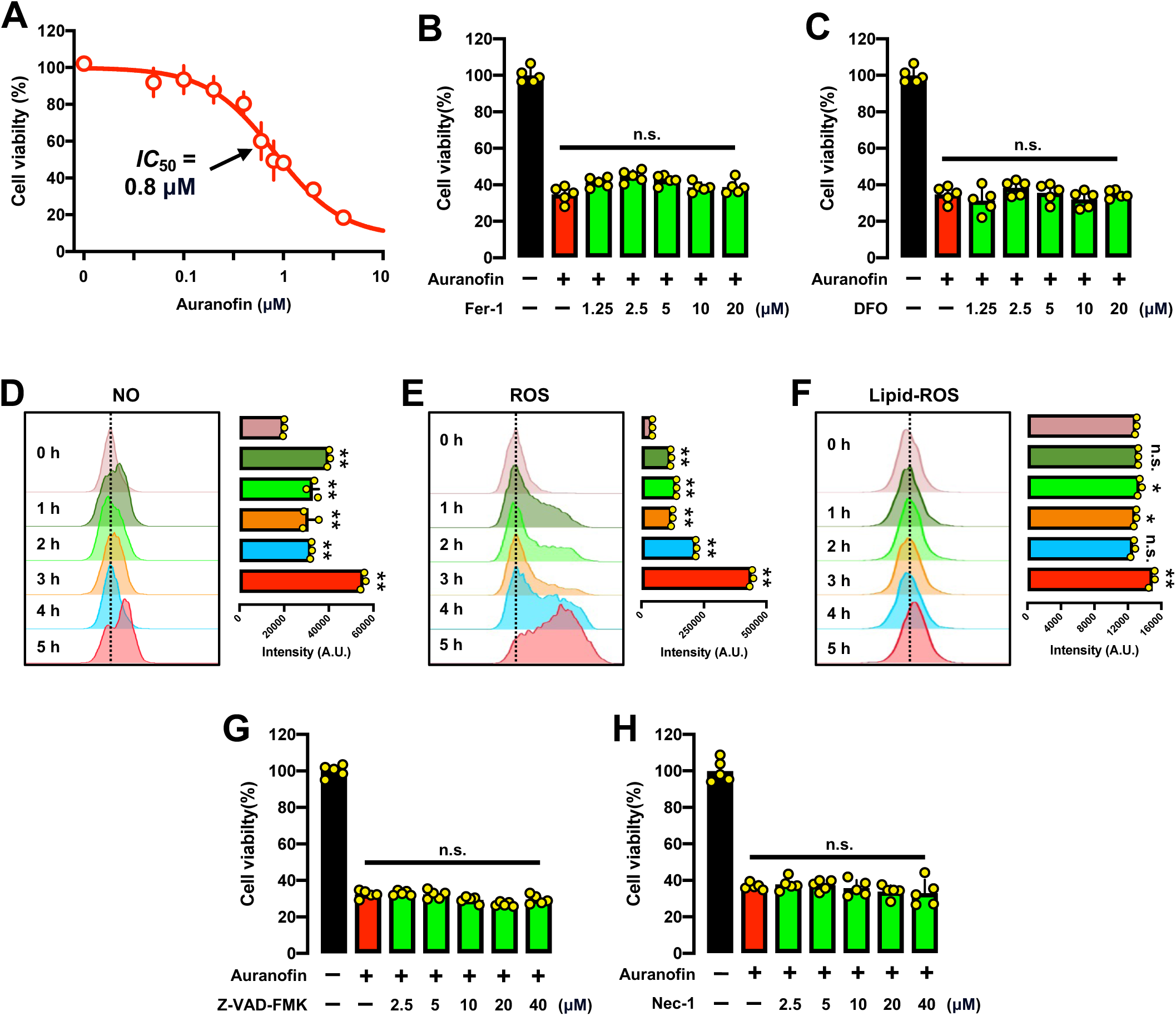
Effect of auranofin on cell viability and cellular levels of NO, ROS and lipid-ROS in HT22 cells. **A.** Dose-dependent effect of auranofin on cell viability (MTT assay, n = 5). **B, C.** Effect of Fer-1 (**B**) and DFO (**C**) on auranofin-induced cytotoxicity. The cells were treated with auranofin (1 µM) ± Fer-1 or DFO at indicated concentrations for 24 h, and change in cell viability was determined by MTT assay (n = 5). **D, E, F.** Time-dependent accumulation of cellular NO (**D**), ROS (**E**) and lipid-ROS (**F**) in auranofin-treated cells (analytical flow cytometry). The quantitative intensity values are shown in the respective right panels (n = 3). **G. H.** Effect of z-VAD-FMK (**G**) and Nec-1 (**H**) on auranofin-induced cytotoxicity. The cells were treated with auranofin (1 µM) ± z-VAD-FMK or Nec-1 at indicated concentrations for 24 h, and change in cell viability was determined by MTT assay (n = 5). Each value represents the mean ± S.D. (\**P* < 0.05; \*\**P* < 0.01; n.s., not significant).

Based on the above observations, we also investigated the effect of jointly treating HT22 cells with auranofin + BSO, which is a GSH synthase inhibitor [40, 41]. It is expected that BSO will facilitate the induction of GSH depletion-associated ferroptosis when they were jointly treated with auranofin. In this experimental design, BSO was used at a concentration (such as 6 µM) that did not cause significant cytotoxicity when present alone (**Fig. 9A**). We found that joint treatment of HT22 cells with very low concentrations of auranofin + 6 μM BSO caused a concentration-dependent reduction in cell viability (**Fig. 9B**). Interestingly, the GPX4 protein level was also decreased in the cells jointly treated with auranofin + BSO (**Fig. 9C**). Notably, a similar decrease in GPX4 protein level was observed in cells treated with RSL3 alone (**Fig. 7D**). The cytotoxicity of auranofin + BSO was partially rescued by Fer-1 (**Fig. 9D**). Cystamine, an inhibitor of PDI which can effectively prevent RSL3-induced ferroptosis (**Fig. 5H**), also exerted a partial protection against the cell death jointly induced by auranofin and BSO (**Fig. 9E**).

**Figure 9.**
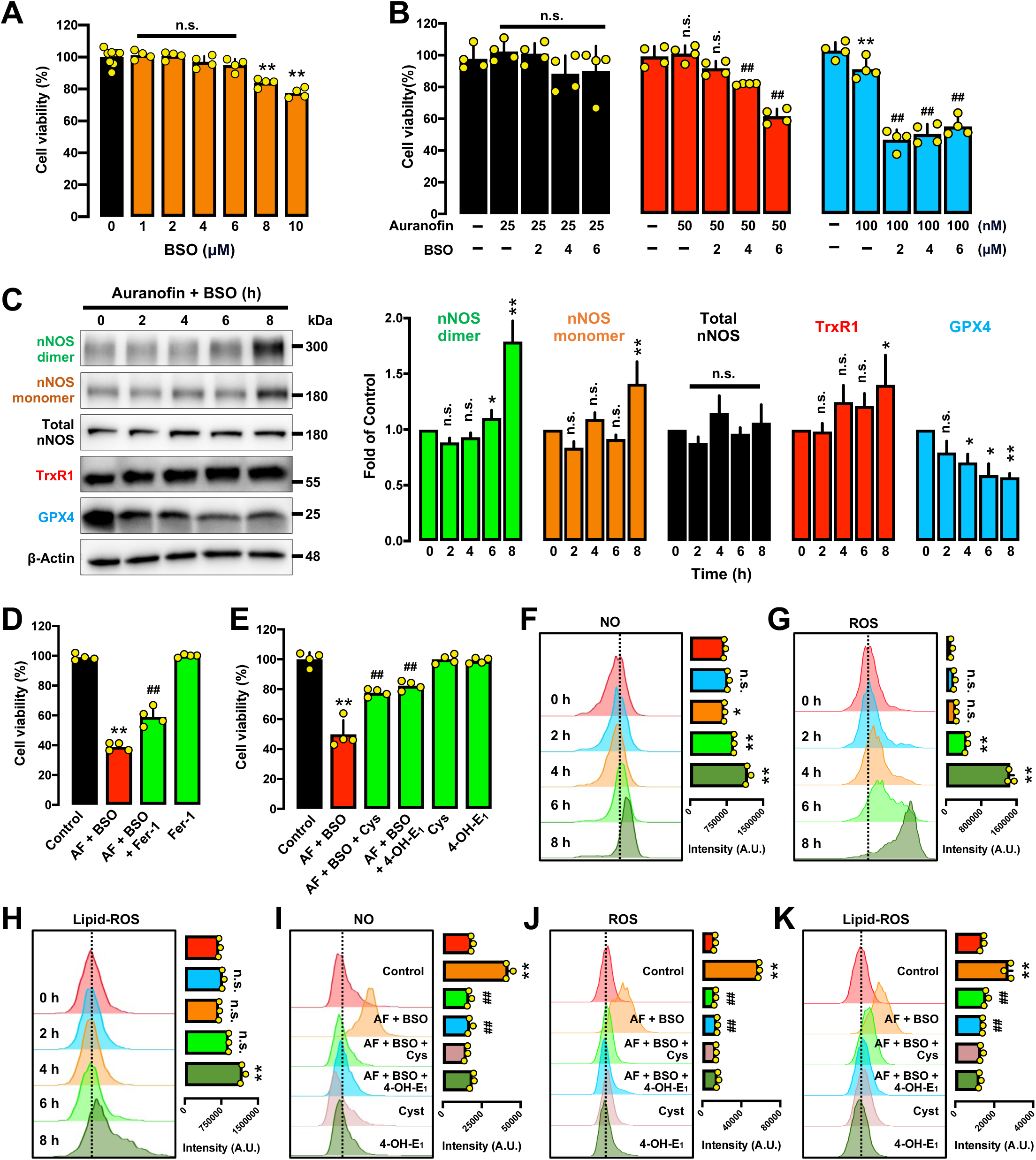
PDI mediates the cytotoxicity jointly induced by auranofin and BSO in HT22 cells. **A.** Dose-dependent effect of BSO on cell viability (MTT assay, n = 4). **B.** Dose-dependent effect of auranofin ± BSO on cell viability. The cells were jointly treated with auranofin (25, 50 and 100 nM) ± BSO (2, 4 and 6 µM) for 24 h, and then cell viability was determined by MTT assay (n = 4). **C.** Protein levels of the monomer and dimer nNOS, total nNOS, TrxR1 and GPX4 in cells jointly treated with auranofin (50 nM) + BSO (6 μM) for different time intervals as indicated. The relative protein levels are shown in the right panels. The relative intensity of protein bands was normalized to β-actin, and calculated relative to the vehicle (control) group. **D.** Protective effect of Fer-1 against cytotoxicity jointly induced by auranofin and BSO. The cells were treated with auranofin (50 nM) + BSO (6 μM) alone or in combination with Fer-1 (1 µM) for 24 h, and cell viability was determined by MTT assay (n = 4). **E.** Protective effect of cystamine and 4-OH-E_1_ against cell death jointly induced by auranofin + BSO. The cells were treated with auranofin (50 nM) and BSO (6 μM) alone or in combination with cystamine (12.5 µM) or 4-OH-E_1_ (2 µM) for 24 h, and cell viability was determined by MTT assay (n = 4). **F, G, H.** Time-dependent accumulation of cellular NO (**F**), ROS (**G**) and lipid-ROS (**H**) in cells jointly treated with auranofin and BSO (analytical flow cytometry). The quantitative intensity values are shown in the respective right panels (n = 3). **I, J, K.** Abrogation by cystamine and 4-OH-E_1_ of auranofin + BSO-induced accumulation of cellular NO (**I**), ROS (**J**) and lipid-ROS (**K**). Cells were treated with auranofin (50 nM) and BSO (6 μM) alone or in combination with cystamine (12.5 µM) or 4-OH-E_1_ (2 µM) for 6 h, and then stained for NO, ROS and lipid-ROS (analytical flow cytometry). The quantitative intensity values are shown in the respective right panels (n = 3). Each value represents the mean ± S.D. (* *P* < 0.05; ** or ^##^ *P* < 0.01; n.s., not significant).

Similarly, 4-OH-E_1_, which can effectively rescue RSL3-induced cell death (**Fig. 6E**), also exerted a partial protection against the cell death jointly induced by auranofin and BSO (**Fig. 9E**). All these observations suggest that when very low concentrations of auranofin (which are close to its *IC*_50_ for TrxR1 inhibition) were used under conditions where cellular GSH is depleted, the cell death induced by auranofin is largely ferroptosis, and as expected, the cell death can be partially prevented by both ferroptosis inhibitors and PDI inhibitors.

To provide further support for the above observations, we also analyzed changes in cellular NO, ROS and lipid-ROS accumulation and NOS dimerization following treatment of cells with very low concentrations of auranofin + BSO. We found that under these joint treatment conditions, there was a time-dependent accumulation of cellular NO (**Fig. 9F**), ROS (**Fig. 9G**) and lipid-ROS (**Fig. 9H**). In addition, the dimeric nNOS (detected as a marker for NOS dimerization) was also significantly increased in a time-dependent manner (**Fig. 9C**). Joint treatment of these cells with cystamine partially abrogated the accumulation of NO (**Fig. 9I**), ROS (**Fig. 9J**) and lipid-ROS (**Fig. 9K**) induced by treatment of auranofin + BSO. Similar observations on the abrogation of NO, ROS and lipid-ROS accumulation were also made when the HT22 cells were jointly treated with 4-OH-E_1_ ± auranofin and BSO (**Fig. 9I-9K**).

## DISCUSSION

RSL3 is a prototypical ferroptosis inducer, and its ferroptosis-inducing ability is thought to be associated with inhibition of GPX4 [4, 42, 43]. However, a recent study showed that in addition to GPX4 inhibition, RSL3 can also inhibit the enzymatic activity of TrxR1 [5], which would shift the pool of thioredoxins to their oxidized forms. PDI is a member of thioredoxin superfamily, and we hypothesize that RSL3 might, through its inhibition of TrxR1’s enzymatic activity [5], keep more cellular PDI proteins in the oxidized form, thereby facilitating RSL3-induced ferroptosis through catalyzing NOS dimerization, followed by cellular ROS and lipid-ROS accumulation, and ultimately ferroptotic cell death (summarized in **Fig. 10**). As discussed below, large amount of experimental data is provided in this study which jointly offers strong support for this hypothesis.

**Figure 10.**
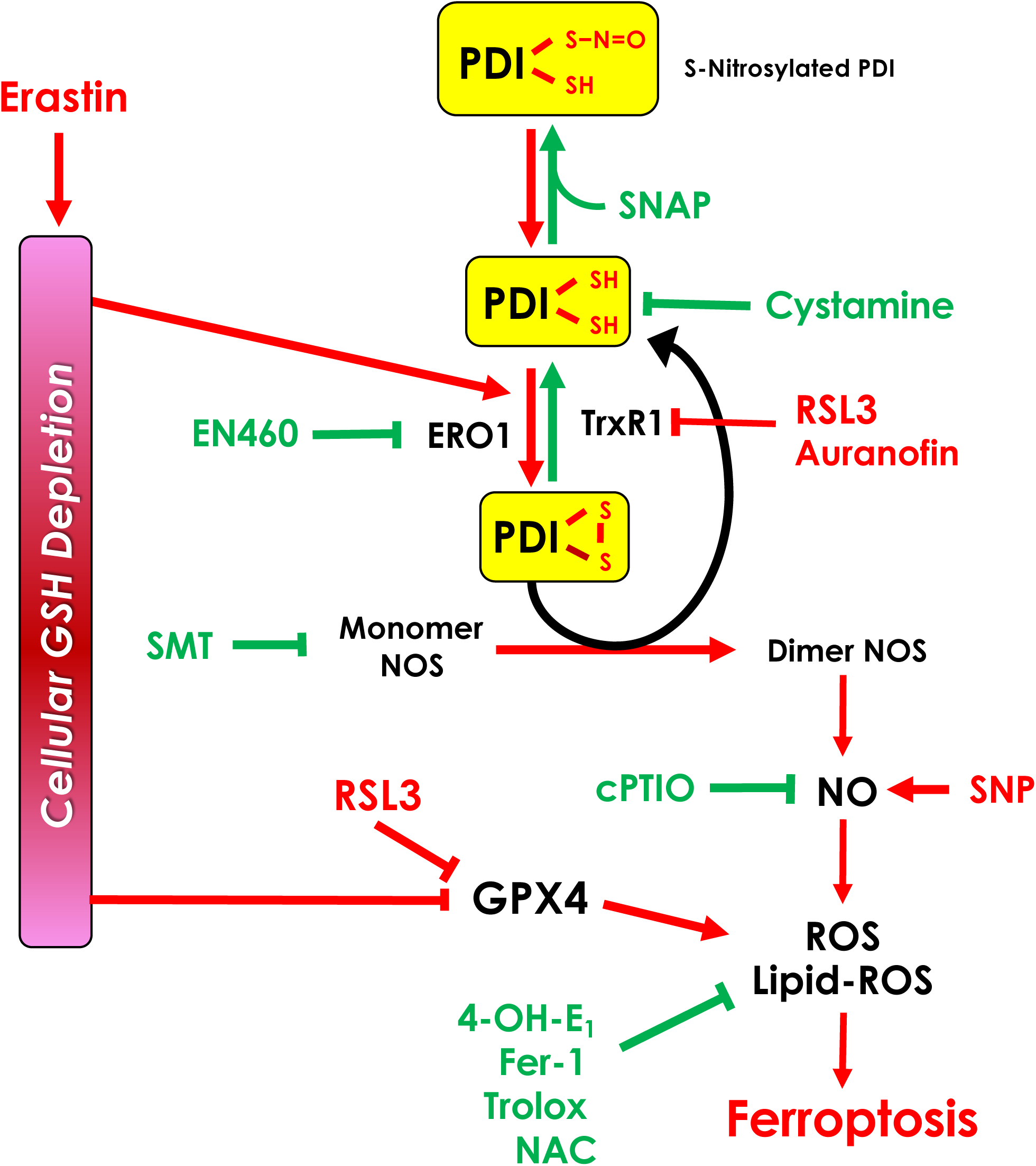
Schematic illustration of the mechanism by which PDI plays an important contributing role in mediating RSL3-induced ferroptotic cell death in HT22 cells. It is proposed that PDI plays an important role in mediating RSL3-induced ferroptotic cell death. PDI is activated in RSL3-treated cells resulting from its inhibition of TrxR1, which subsequently leads to PDI-mediated NOS dimerization (activation) and accumulation of cellular NO, ROS and lipid-ROS, and ultimately ferroptotic cell death.

We find that during the development of RSL3-induced ferroptosis in HT22 cells, NO accumulation is an early event which subsequently leads to the formation and accumulation of ROS and lipid-ROS, contributing to the development of ferroptotic cell death. The supporting evidence includes: First, we find that RSL3 can induce the accumulation of both NO and ROS/lipid-ROS in a time- and dose-dependent manner, although there appears to be no clear time differential in their buildup (the reason for this is provided later). The observations made with SNP, which can release NO [21], offer support for this suggestion. We find that SNP can quickly lead to accumulation of cellular ROS and lipid-ROS when present alone in HT22 cells, and it also dramatically sensitizes the cells to RSL3-induced ROS/lipid-ROS accumulation and ferroptosis. Here it is of note that our earlier study showed that during erastin-induced ferroptosis, NO accumulation occurred earlier than did ROS/lipid-ROS accumulation in HT22 cells [12].

In addition, when the HT22 cells were jointly treated with erastin + SNP, the NO accumulation also occurred much earlier (due to the rapid release of NO from SNP), and in parallel manner, the ROS/lipid-ROS accumulation also occurred much earlier, lagging shortly behind NO accumulation [12]. These experimental observations jointly suggest that during chemically-induced ferroptosis, NO accumulation occurs first, which then leads to ROS/lipid-ROS accumulation. The second line of evidence comes from the study of NO scavengers and NOS inhibitors. We find that cPTIO, a NO scavenger, can abrogate RSL3-induced NO accumulation, and this effect is accompanied by abrogation of ROS/lipid-ROS accumulation and ferroptosis. In addition, SMT, an iNOS inhibitor, exerts a strong protection against RSL3-induced NO accumulation, which is accompanied by abrogation of ROS/lipid-ROS accumulation and ferroptotic cell death. Together, these observations suggest that RSL3-induced NO accumulation contributes to the subsequent accumulation of ROS and lipid-ROS, and ultimately the induction of ferroptotic cell death.

Although RSL3-induced NO accumulation leads to cellular ROS/lipid-ROS accumulation in HT22 cells, it is interesting that RSL3-induced buildup of cellular NO and cellular ROS (which includes lipid-ROS) do not display a distinguishable time difference; in fact, they occur nearly at the same time. This unexpected experimental observation likely is attributable to the fact that RSL3 can increase both NO and lipid- ROS accumulation at the same time. RSL3 increases NO biosynthesis through its direct inhibition of TrxR1, which then leads to PDI activation and NOS dimerization; on the other hand, RSL3 also directly increases cellular lipid-ROS accumulation (independent of cellular NO accumulation) due to its inhibition of GPX4. Therefore, the buildup of cellular lipid-ROS (which is part of the cellular ROS) in RSL3-treated cells is the result of both GPX4 inhibition (which likely occurs first) and NO accumulation (which likely occurs at a slightly later time). Because of these reasons, the accumulation of cellular NO and ROS/lipid-ROS occurs rapidly and almost at the same time after exposure to RSL3.

The dimeric NOS is catalytically active and catalyzes NO formation [12–16]. In this study, we show that RSL3 exposure increases the formation of nNOS and iNOS dimers in a time- and dose-dependent manner in HT22 cells. The results from iNOS and nNOS inhibitor experiments support the idea that both iNOS and nNOS are involved in RSL3- induced ferroptosis, but the results with iNOS and nNOS knockdown experiments are inclusive. It is shown that iNOS knockdown is associated with a partial abrogation of RSL3’s cytotoxicity, but nNOS knockdown fails to exert a significant protection (**Fig. 5**). Two potential reasons may underlie the inability to observe a significant protective effect with nNOS knockdown: First, it is likely that the relative contribution of cellular nNOS (which is not induced following RSL3 exposure) to the cellular NO pool is markedly smaller than the contribution of cellular iNOS (which is strongly-induced by RSL3). The other reason might be due to the low effectiveness of a sustained nNOS protein knockdown in live HT22 cells. Together, it is apparent that iNOS clearly plays a more important role than nNOS in mediating RSL3-induced NO accumulation and ferroptosis in this neuronal cell culture model.

By jointly using chemical inhibitors or PDI knockdown, strong evidence is provided in this study to show that PDI is involved in catalyzing NOS dimer formation in RSL3- treated HT22 cells. For instance, joint treatment of HT22 cells with cystamine (a PDI inhibitor) strongly reduces RSL3-induced dimerization of both nNOS and iNOS, the accumulation of NO, and ROS/lipid-ROS, and the oxidative cytotoxicity. Another example involves the *S-*nitrosylation of the cysteine residues in PDI’s active site, which inactivates PDI’s enzyme activity [44, 45]. Our earlier study showed that PDI in untreated HT22 cells was mostly present in the *S*-nitrosylated form, but following glutamate treatment (which induces cellular GSH depletion), PDI underwent *S*- denitrosylation [25]. We find in this study that RSL3-induced NO, ROS and lipid-ROS accumulation is abrogated by SNAP (which causes PDI *S*-nitrosylation), and this effect associated with a protection against cell death. Together, these observations offer support for the notion that PDI plays an important role in catalyzing NOS dimerization in RSL3-treated HT22 cells, which then leads to accumulation of NO, ROS and lipid-ROS, and ultimately ferroptotic cell death.

ERO1 is an enzyme that oxidizes PDI by forming disulfide bonds in its active site [46]. Our recent work showed that EN460 (an inhibitor of ERO1) has a strong protective effect against erastin-induced ferroptosis in HT22 cells [12]. However, EN460 only shows a very weak protective effect against RSL3-induced ferroptosis as observed in this study (**Fig. S3A –S3D**). This difference is because RSL3 is an inhibitor of TrxR1, which catalyzes the reduction of the disulfide bonds in PDI’s active site. When the cellular TrxR1 activity is inhibited by RSL3, the majority of PDI will be in the oxidized form; as such, the presence of ERO1 will not add significantly more oxidized PDI (because not much reduced PDI is left as substrate for ERO1). This is why it will be difficult to display a strong protective effect of EN460 when TrxR1 is already inhibited by RSL3. This lack of a significant protective effect of EN460, in fact, also supports the important role of PDI in driving RSL3-mediated ferroptosis.

Based on the findings of this study, it is evident that RSL3-induced ferroptosis involves two important mechanistic components: one is mediated by the activation of PDI (resulting from TrxR1 inhibition by RSL3), and other component is the inhibition of the GPX4-mediated protection against lipid-ROS. Because of the combination of these two effects, a PDI inhibitor which does not have a direct antioxidant activity is expected to be less efficacious in protecting against RSL3-induced ferroptosis compared to a PDI inhibitor which also has a strong antioxidant activity. For instance, 4-OH-E_1_, which was recently identified as an effective PDI inhibitor [33], is found to have a strong protective effect against RSL3-induced ferroptotic cell death. The protective effect of 4-OH-E_1_ against RSL3-induced cell death is more potent than its protection against erastin- induced cell death in some of the cell lines (unpublished data). This observation suggests that the direct antioxidant activity of 4-OH-E_1_ against lipid-ROS also contributes, to some degrees, to its cytoprotective effect. This explanation is also in line with the observation with cystamine, which can inhibit PDI function but does not have any antioxidant property. Cystamine exerts a less efficacious protection against RSL3- induced ferroptosis in HT22 cells compared to 4-OH-E_1_; however, both cystamine and 4- OH-E_1_ exert a similarly strong protection (90l7100% protection) against erastin- induced ferroptosis in the same cells (data not shown).

In this study, we find that the TrxR1’s enzyme activity can be directly and potently inhibited by auranofin (with an apparent *IC*_50_ of approximately 1.1 nM), which is slightly lower than the reported *IC*_50_ (∼20 nM) for human TrxR1 [47]. Notably, it is observed in this study that when the *in vitro* reaction time is prolonged, the reduction in product formation becomes progressively greater, and there is a sustained reduction in product formation (reflecting a sustained TrxR1 inhibition). This observation suggests that auranofin inhibits TrxR1’s catalytic activity resulting from its covalent modification of the enzyme. Indeed, earlier studies have reported that auranofin can inactivate TrxR1 by forming covalent diselenide bridges with its Sec498 residue [39, 48].

We find that the potency of RSL3 to inhibit TrxR1 *in vitro* is markedly lower than that of auranofin. RSL3’s apparent *IC*_50_ (approximately 4 μM) detected in this study is in line with the results of an earlier study [5]. Based on the similar patterns of TrxR1 inhibition by RSL3 and auranofin observed in this study, it is suggested that RSL3 may also inhibit TrxR1 in a covalent manner. This suggestion is partially supported by the observation made in HT22 cells showing a high-potency inhibition of the TrxR1 activity by RSL3. It should be noted that the apparent discrepancy between the high-potency RSL3 inhibition in intact cells and the low-potency inhibition of the TrxR1’s catalytic activity in an *in vitro* enzymatic assay may suggest that RSL3 cannot readily access the catalytic site of TrxR1 in an aqueous reaction system *in vitro*. A similar phenomenon was also observed in an earlier study with the flavonoid-type compounds: while these compounds can strongly activate the cyclooxygenase’s catalytic activity in live cells in culture when present at low nM concentrations, they needed to be present at 100l7200 μM concentrations in order to exert a similar activation of the pure enzymes in the *in vitro* enzyme assays [49].

In conclusion, the results of our present study demonstrate that PDI plays a critical role in mediating RSL3-induced ferroptotic cell death in HT22 cells by catalyzing the dimerization of both nNOS and iNOS, followed by cellular accumulation of NO, ROS and lipid-ROS, and ultimately ferroptotic cell death (summarized in **Fig. 10**). The discovery of an important role of PDI in mediating chemically-induced ferroptosis suggests that PDI might serve as a potential drug target for protection against ferroptosis-related neuronal degeneration.

## Data Availability Statement

The authors confirm that the data supporting the findings of this study are available within the article and its supplementary materials.

## Abbreviations

PDI: protein disulfide isomerase
nNOS: neuronal nitric oxide synthase
iNOS: inducible nitric oxide synthase
NO: nitric oxide
ROS: reactive oxygen species
TrxR1: thioredoxin reductase 1

## FIGURE LEGENDS

**Supplementary Figure S1. Additional data comparing the time-dependent accumulation of cellular NO (A) and ROS (B) in RSL3-treated cells (analytical flow cytometry).** All the experimental conditions were the same as described for Fig. 1G and 1H, except that a 3-h time point was added in this experiment. Each value is the mean ± S.D. (n = 3).

Each value represents the mean ± S.D. (**P < 0.01; n.s., not significant).

**Supplementary Figure S2. Effect of TrxR1 ± PDI knockdown by siRNAs on RSL3- induced ferroptosis in HT22 cells.**

**A.** Effect of TrxR1 knockdown by TrxR1-siRNAs on the viability of HT22 cells (MTT assay, n = 5). Cells were transfected with the control and TrxR1-specific siRNAs for 24 h, and then the cells were treated with a vehicle (without RLS3) for 24 h. Then the MTT assay was performed, and the absolute absorbance values (after subtracting the background value) are shown.

**B.** Effect of joint knockdown of PDI and TrxR1 by siRNAs on the sensitivity to 100 nM RSL3-induced cell death (MTT assay, n = 5). The viability of vehicle-treated cells in the siCon group or in the siPDI + siTrxR1 group are individually set as 100% when compared with the respective RSL3-treated cells.

Each value represents the mean ± S.D. (** or ^##^ P < 0.01).

**Supplementary Figure S3. Effect of EN460 (a ERO1 inhibitor) on RSL3-induced ferroptosis in HT22 cells.**

**A.** Effect of EN460 on RSL3-induced cytotoxicity following treatment of cells with RSL3 (100 nM) ± EN460 (5, 10 and 20 µM) for 24 h (MTT assay, n = 4).

**B, C, D.** Effect of EN460 on RSL3-induced accumulation of cellular NO (B), ROS (C) and lipid-ROS (**D**). Cells were treated with RSL3 (100 nM) ± EN460 (20 µM) for 6 h, and then stained for NO, ROS and lipid-ROS (**B, C** for fluorescence microscopy; scale bar = 65 µm; **D** for confocal microscopy, scale bar = 10 µm).

Each value represents the mean ± S.D. (** or ^##^ P < 0.01; n.s., not significant).

