## Supplementary figures and images for "Mechanism of RSL3-Induced Ferroptotic Cell Death in HT22 Cells: Crucial Role of Protein Disulfide Isomerase"

### Supplemental Figure S1-3

#### Supplementary Figure S1

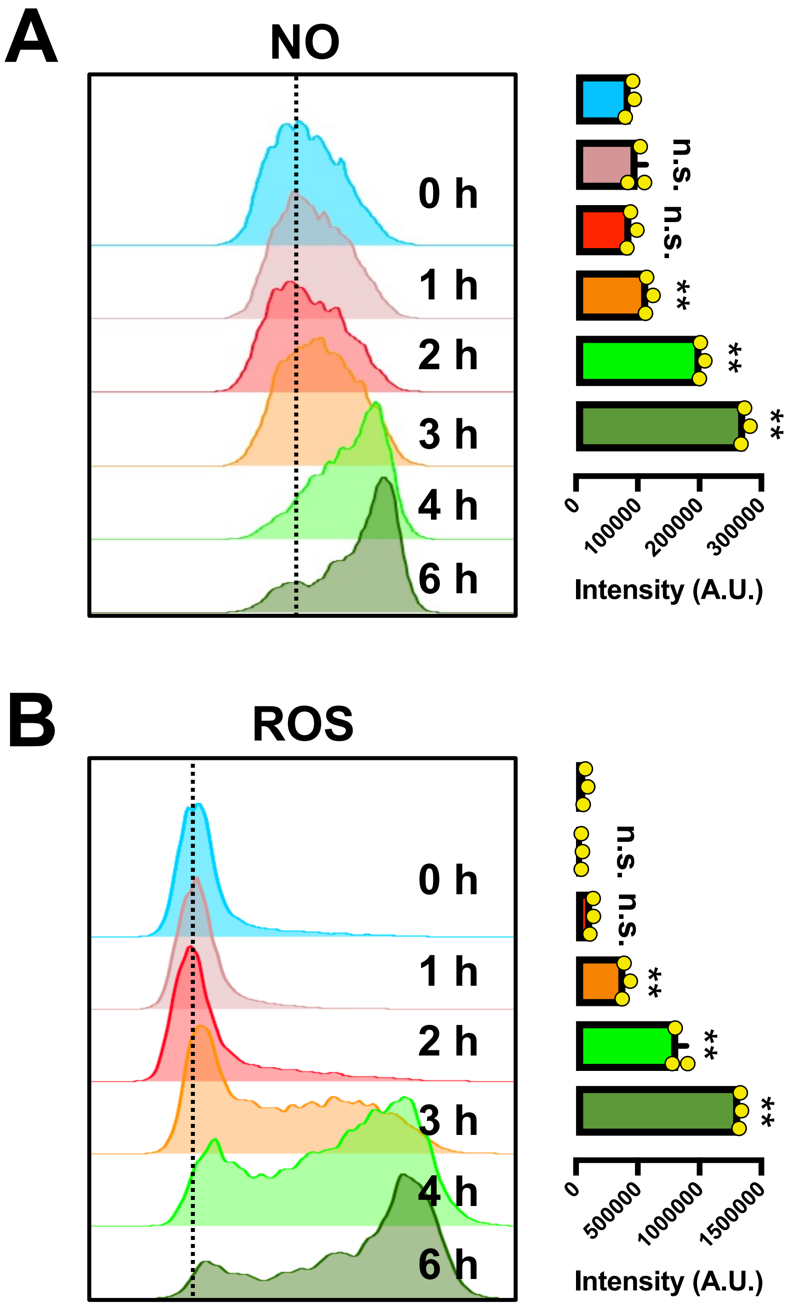

##### Supplementary Figure S2

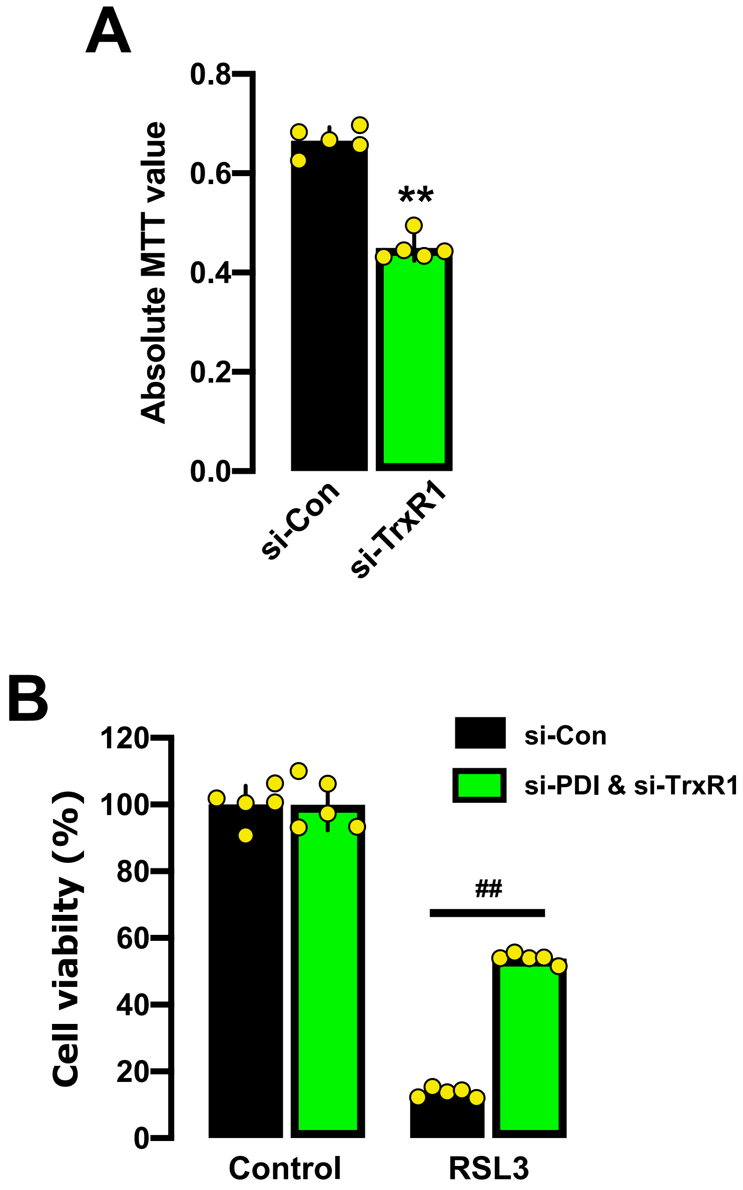

#### Supplementary Figure S3

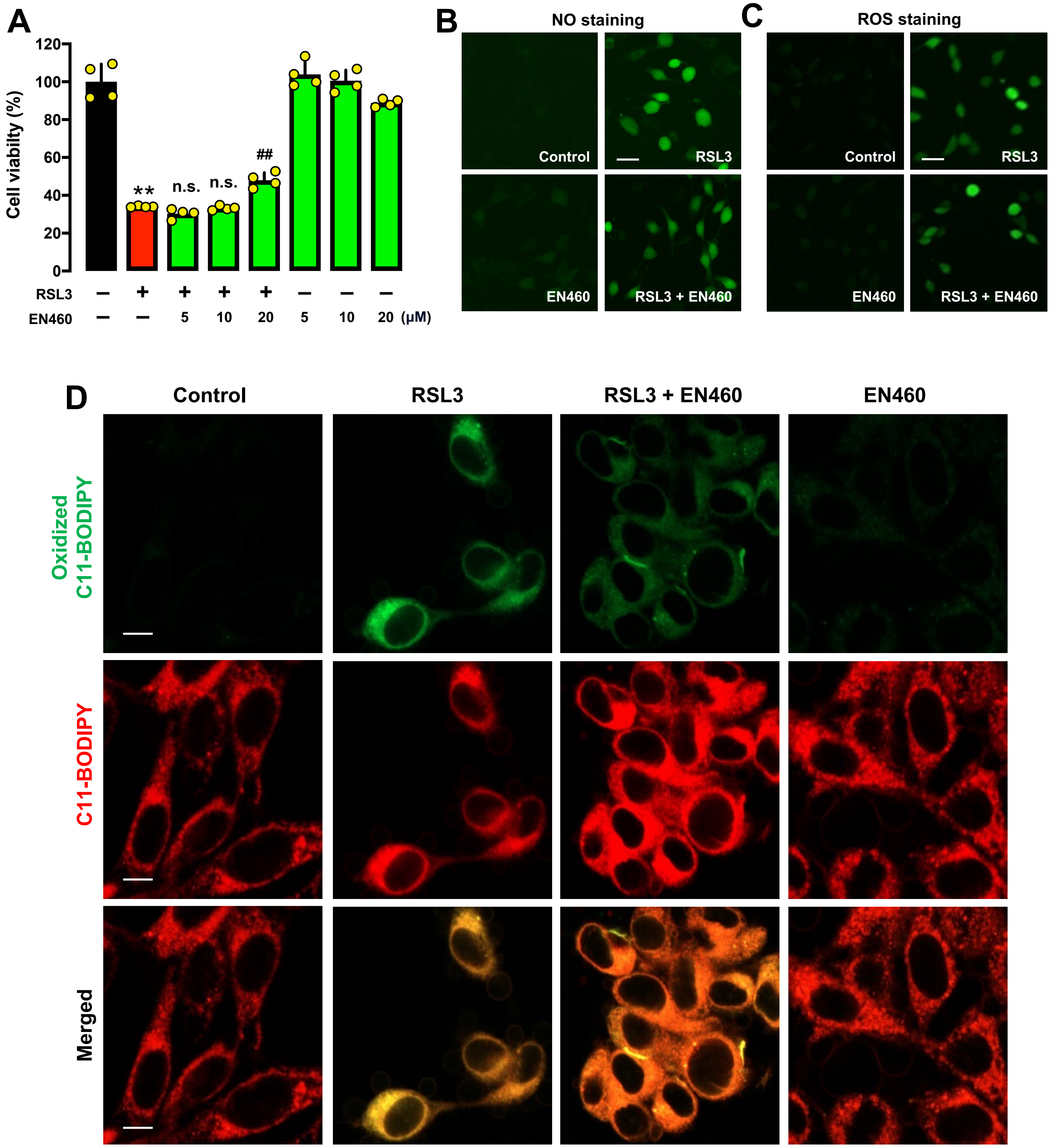
